# A survey of human cancer-germline genes : linking X chromosome localization, DNA methylation and sex-biased expression in early embryos

**DOI:** 10.1101/2025.05.19.654804

**Authors:** Axelle Loriot, Julie Devis, Laurent Gatto, Charles De Smet

## Abstract

Human cancer-germline (CG) genes are a group of testis-specific genes that become aberrantly activated in various tumors. Ongoing studies aim to understand their functions in order to evaluate their potential as anti-cancer therapeutic targets. Evidence suggests the existence of subcategories of CG genes, depending on location on autosomal or sex chromosomes, reliance on DNA methylation for transcriptional regulation, and profile of expression during gametogenesis and early embryogenesis. To clarify this issue, we developed CTexploreR, a R/Bioconductor package that integrates an up-to-date reference list of human CG genes (n=146) with multiple bulk and single-cell methylomic and transcriptomic datasets. Based on promoter methylation profiles and responsiveness to a DNA methylation inhibitor, 74% of the CG genes were classified as DNA methylation dependent (Methdep). Intriguingly, most X-linked CG genes (69/70) fell into this category, thereby implicating DNA methylation dependency in the well-documented over-representation of testis-specific genes on the X chromosome. We further observed that, whereas X-linked Methdep CG genes become demethylated and activated in pre-spermatogonia in the fetal testis, most of them resist DNA demethylation in female germ cells and remain therefore silent in fetal and adult oocytes. Importantly, a number of X-linked Methdep CG genes (e.g. *FMR1NB, GAGE2A, MAGEB2/C2, PAGE2, VCX3A/B*) maintained this maternal-specific imprinting after fertilization, and were expressed exclusively in female preimplantation embryos, which inherit a paternal X chromosome. Together, our study using the CTexploreR package has allowed us to show that X-linked CG genes undergo transient maternal imprinting and contribute therefore to transcriptional sexual dimorphism in early embryos.

**Author Summary:** Cancer-germline (CG) genes include a set of genes that are normally active only in the testis but become aberrantly switched on in different types of tumor, making them potential targets for new anti-cancer treatments. We developed a new analytical tool called CTexploreR, to study these genes, and observed that a large subset of CG genes reside on the X chromosome and use DNA methylation as a mechanism of repression in non-testicular tissues. In fetal testis, these genes lose methylation and become activated in early spermatogenic cells, while in female ovaries they stay methylated and remain silent. Notably, we demonstrate that several CG genes keep this methylation pattern after fertilization, and are therefore expressed in female but not male embryos, which inherit only one X chromosome of maternal origin. CG genes carrying this transient maternal imprinting appear therefore as main contributors of sex-biased mRNA expression in preimplantation embryos. Our Jindings therefore open up new Jields of investigation into the functions of CG genes, the sexual dimorphism of early embryos, and the intergenerational transmission of epigenetic imprints.

## Introduction

Cancer-Germline (CG) genes, also called Cancer-Testis (CT) genes, are a group of genes that are normally expressed exclusively in testicular germ cells, but become aberrantly activated in a significant proportion of tumors of different histological origins. Genes with this particular expression profile were initially identified on the basis of their ability to produce tumor-specific antigens recognized by cytolytic T lymphocytes [1]. The immunogenicity of CG gene products results from their restricted expression in germ cells, which do not express antigen-presenting major histocompatibility complexes (MHC) [2]. In contrast, activation of CG genes in tumors of somatic origin gives rise to antigenic peptides that are presented at the cell surface and can be recognized as non-self by the immune system. CG genes represent therefore ideal targets for anti-cancer vaccines [3,4]. Their unique expression profile, however, extends their clinical potential beyond cancer immunotherapy. It is anticipated indeed that they may be useful as cancer biomarkers [5], and may represent appealing targets for the development of anti-cancer therapies with limited side effects [5]. In this regard, studies are underway to elucidate the still poorly understood cellular functions of CG genes, in order to determine whether they contribute to oncogenic pathways [6–8].

Since the initial discovery of CG genes, many others have been identified through either cloning of tumor antigen-encoding genes or transcriptional profiling. Isolated CG genes were found to share several features. First, a majority of CG genes reside on the X chromosome [9], and this is believed to result from evolutionary constraints for genes with testis-specific functions. The “sexual antagonism” theory states that genes conferring reproductive advantages to males, but not females, become enriched on the X chromosome during evolution [10]. It remains however unclear whether CG genes exert sex-related functions. A further observation is that many CG genes residing on the X chromosome belong to gene families, and have a recent evolutionary origin, some being specific to the human species [11].

A striking feature of CG genes is that many of them use DNA methylation, a chemical modification of cytosines in CpG sequences, as a primary mechanism of transcriptional repression in somatic tissues [12,13]. Aberrant activation of these genes in tumors can thus be explained by the process of global DNA demethylation that often accompanies tumorigenesis [14–16]. However, the role of DNA methylation has not been investigated for all CG genes, and it is therefore uncertain if this mechanism of epigenetic regulation applies to all or only part of them.

In the adult testis, expression of CG genes was observed at various stages of spermatogenesis. Surprisingly, CG genes located on the X chromosome display preferential expression in pre-meiotic stages, particularly in spermatogonia [17]. Instead, CG genes residing on a chromosome other than the X are often expressed at later stages of germ cell differentiation, including spermatocytes and spermatids. The reason for this chromosome location-dependent expression pattern remains unexplained.

A few CG genes were shown to be also expressed in female germ cells [18,19]. It is not clear, however, whether this can be generalized to all CG genes. Investigating gene expression in the female germline is not an easy task, because pre-meiotic germ cells are only present in the fetal ovary, and oocytes represent only a small proportion of the cells that make up the ovarian tissue in the adult.

Finally, several studies reported expression of CG genes in the embryo [20,21]. Uncertainty remains, however, as to the embryonic stages where this expression takes place, as well as the extent to which embryonic expression applies to most or only few CG genes.

Considering the issues described above, it seems that CG genes may actually fall into different categories, depending on chromosomal location, transcriptional regulation and expression profile. Establishing such a sub-classification would however require access to a database integrating the various characteristics of CG genes. Currently, the reference database is the CTdatabase [22], a literature-based repository that was published in 2009. The CTdatabase references 276 CG genes, but it is no longer updated. A recent re-evaluation of their expression in normal tissues using omics data revealed that some genes in this database do not exhibit the expected tissue-specific expression [6]. Another limitation of the CTdatabase is that it is not in an easily importable format, and that some genes are not named properly, altogether resulting in poor interoperability for downstream analyses. More recent omics studies gave rise to other lists of CG genes [23–26]. These lists differ however substantially between each other, mainly because they use various criteria to define CG genes. Moreover, many of the lists are provided as supplemental data files rather than searchable databases. Lastly, none of these studies explores the involvement of DNA methylation in the regulation of individual CG genes.

Here we present CTexploreR [27], a R/Bioconductor package that integrates an up-to-date reference list of CG genes with multiple omics databases, including methylomes and transcriptomes of normal and cancerous tissues and cell lines, as well as single-cell RNA-Seq and Bisulfite-Seq data of male and female gonads, and of early embryos. Using CTexploreR [27], we have explored the existence of subcategories of CG genes, in relation to DNA methylation dependency, chromosomal localization, and expression profile during gametogenesis and embryogenesis.

## Results

### CTexploreR package

As a first step, we used publicly available omics data to implement a reliable list of CG genes (**Fig. 1A**). RNAseq data of healthy tissues from the Genotype-Tissue Expression (GTEx) database [28] were examined, and genes with expression strictly limited to testis were selected. We noticed, however, that some well-known CG genes, (e.g. genes of the *MAGE, SSX, CT45A*, and *GAGE* families) were undetectable in the GTEx testis samples. This is likely because of the high sequence similarities between members in these gene families, which cause multi-mapping of mRNA reads (i.e. reads aligned to different genomic regions) and consequent elimination in the GTEx RNAseq processing pipeline. To overcome this issue, raw RNAseq data of normal tissues were processed with multi-mapping support. Under these conditions, CG genes that were initially missing became observable (Supplemental **Fig. S1**), and were added to the list of testis-specific genes. We next sought to eliminate genes that show expression in somatic cells of the testis, as well as in discrete cell population of somatic tissues. To this end, the gene list was crossed with the Single Cell Type Atlas classification [29] of the Human Protein Atlas [30] to exclude genes that were alagged as specific of any somatic cell type. This led to a final list of 1493 genes displaying highly restricted expression in testicular germ cells.

**Figure 1.**
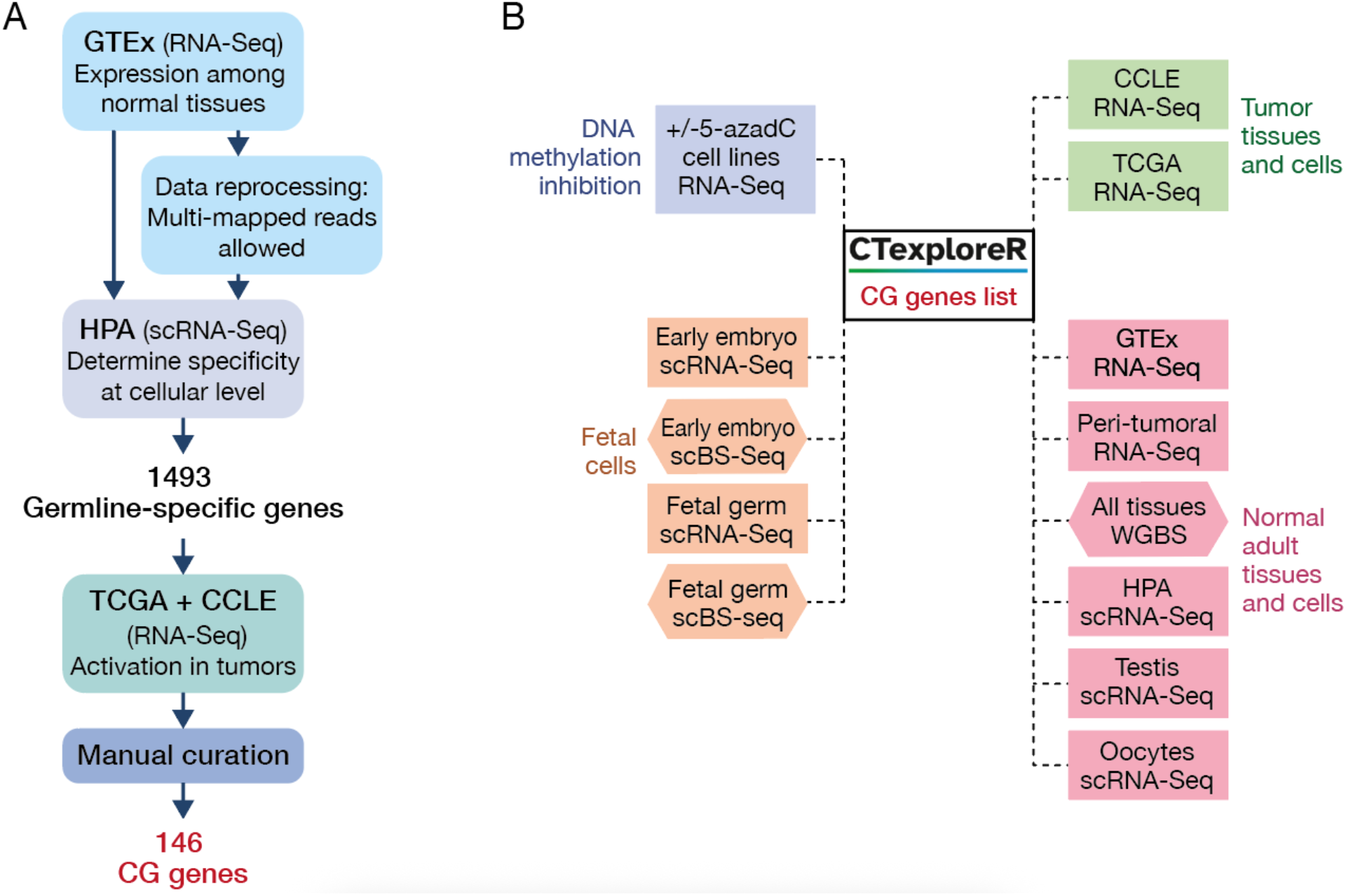
CTexploreR. **(A)** Summary of the work/low applied for selection of CG genes. **(B)** Omics datasets integrated into the package.

Next, selected genes were screened in order to identify those that become activated in tumor cells. To this end, RNAseq data from both The Cancer Genome Atlas [31] (TCGA) and the Cancer Cell Line Encyclopedia [32] (CCLE) were screened, and genes showing transcriptional activation in at least 1% of tumors and cancer cell lines of any histological type were selected. In an additional effort to select genes with high expression specificity, we further eliminated at this stage genes that exceeded a threshold level of expression (TPM > 0.5 in > 25% samples) in normal peri-tumoral tissues (TCGA).

Our workalow led to a final list of 146 CG genes displaying highly specific expression in testicular germ cells and significant activation in tumor cells (supplemental **Table S1**). These genes were transcribed in a table that constitutes the core of the CTexploreR package [27].

We integrated multiple omics datasets into the package, and developed functions that enable to visualize expression and promoter DNA methylation of CG genes in normal and tumoral tissues, at either bulk or single-cell level (**Fig. 1B**).

Of note, we also compiled a list of genes referred to as “CG-preferential” (n=134), which showed an expression that was not strictly limited to testis (testis-preferential), but nevertheless displayed significant up-regulation in different tumors (Supplemental **Table S2**). We will, however, focus our analyses on the 146 highly specific CG genes.

### Comparison of CTexploreR with other CG gene databases

We compared our list of 146 CG genes with those reported in the CTdatabase [22]. Strikingly, 183 out of the 276 genes included the CTdatabase were not present in our list of CG genes (**Fig. 2A**), because they were considered non testis-specific and/or not activated in cancer by CTexploreR. On the other hand, 83 of the CG genes we selected in CTexploreR had not been previously reported in the CTdatabase. As mentioned above, other lists of CG genes have been established more recently based on omics data analyses [23–26], but with no consistency in the selection approaches used. Not surprisingly, the amount of selected CG genes varies greatly from one list to another, and overlaps between studies are usually very low (**Fig. 2B**). Despite this marked heterogeneity, we observed that 98 of the CG genes listed in CTexploreR (67%) are also present in at least one of the other gene lists (**Fig. 2B**). Importantly, 48 genes selected in CTexploreR had never before been classified as cancer-germline. Together, these observations suggest that our list represents a very stringent, yet highly representative, selection of CG genes.

**Figure 2.**
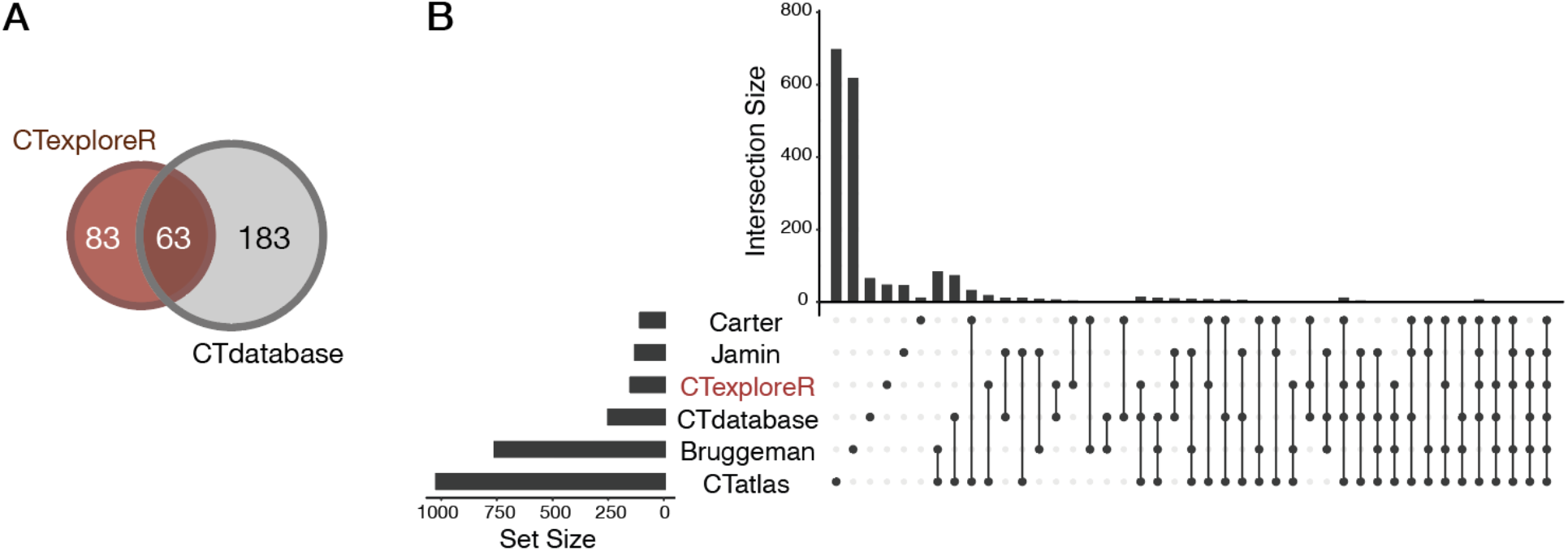
Comparison of the CG gene list of CTexploreR with other cancer-testis gene lists. **(A)** Venn diagram showing overlap between CG genes reported in CTexploreR (brown) and in CTdatabase (grey). Note that the analysis was applied to only 246 genes of the CTdatabase, because 30 out of the 276 originally reported genes were wrongly annotated. **(B)** Upset plot showing all intersections between CTexploreR and /ive other published cancer-testis gene lists, sorted by number of intersections and number of intersecting genes. Solid circles in the matrix indicate lists that are part of the intersection.

### A majority of CG genes exhibit DNA methylation dependency

Having generated an interoperable dataset of CG genes, we first set out to determine which of these rely on promoter DNA methylation as a mechanism of transcriptional regulation. To this end, we first explored whole genome bisulfite sequencing (WGBS) data of normal human tissues, in order to identify CG genes displaying consistent promoter methylation in non-expressing somatic tissues. Additionally, we interrogated RNAseq data of cell lines (n=8) that had been exposed to the DNA methylation inhibitor 5-Aza-2′-deoxycytidine (5-AzadC) [33–36], with the aim to identify CG genes that become induced by the treatment. CG genes that met the two criteria (expected promoter methylation profile, and transcriptional induction by 5-AzadC) were classified as “ methylation dependent” (Methdep). For a few CG genes (n=22, mostly including multi-mapping genes like *MAGE, GAGE, CT45*), only the second criteria was used for classification, because WGBS data were missing. On the basis of our filtration criteria, 108/146 (74%) CG genes qualified as Methdep (**Fig. 3A**). Visualization of the WGBS data confirms that the promoter methylation level of Methdep CG genes is generally high in somatic tissues, and substantially lower in testis (mixture of somatic and germ cells) and sperm (**Fig. 3B**). Promoters of Methdep CG genes also showed intermediate DNA methylation levels in placenta, a temporary organ where the expression of several CG genes has been reported previously [37]. The promoters of non-Methdep CG genes instead showed highly variable levels of methylation, with no marked difference between somatic and germline tissues (**Fig. 3B**). The heatmap representation of RNA-seq data of 5-AzadC-treated cell lines (**Fig. 3C**), shows significant up-regulation of Methdep CG genes, but not non-Methdep CG genes, upon exposure to the DNA methylation inhibitor in at least one cell line. Together, these results suggest that CG genes fall into two categories on the basis of their dependency on DNA methylation for tissue-specific expression: a majority being dependent on their promoter DNA methylation status (Methdep, 74%), and the other part (non-Methdep, 26%) probably relying on other regulatory mechanisms. Because the density of CpGs impacts on the regulatory effects of DNA methylation, we examined the DNA sequences of CG gene promoters. We observed that, compared to non-Methdep CG genes, Methdep CG genes more often contain a promoter with an intermediate density of CpGs (Fisher’s exact test: p-value = 0.0002; **Fig. 3D**). This is consistent with previous observations indicating that promoters with an intermediate density of CpGs are more prone to acquire tissue-specific methylation patterns [38].

**Figure 3.**
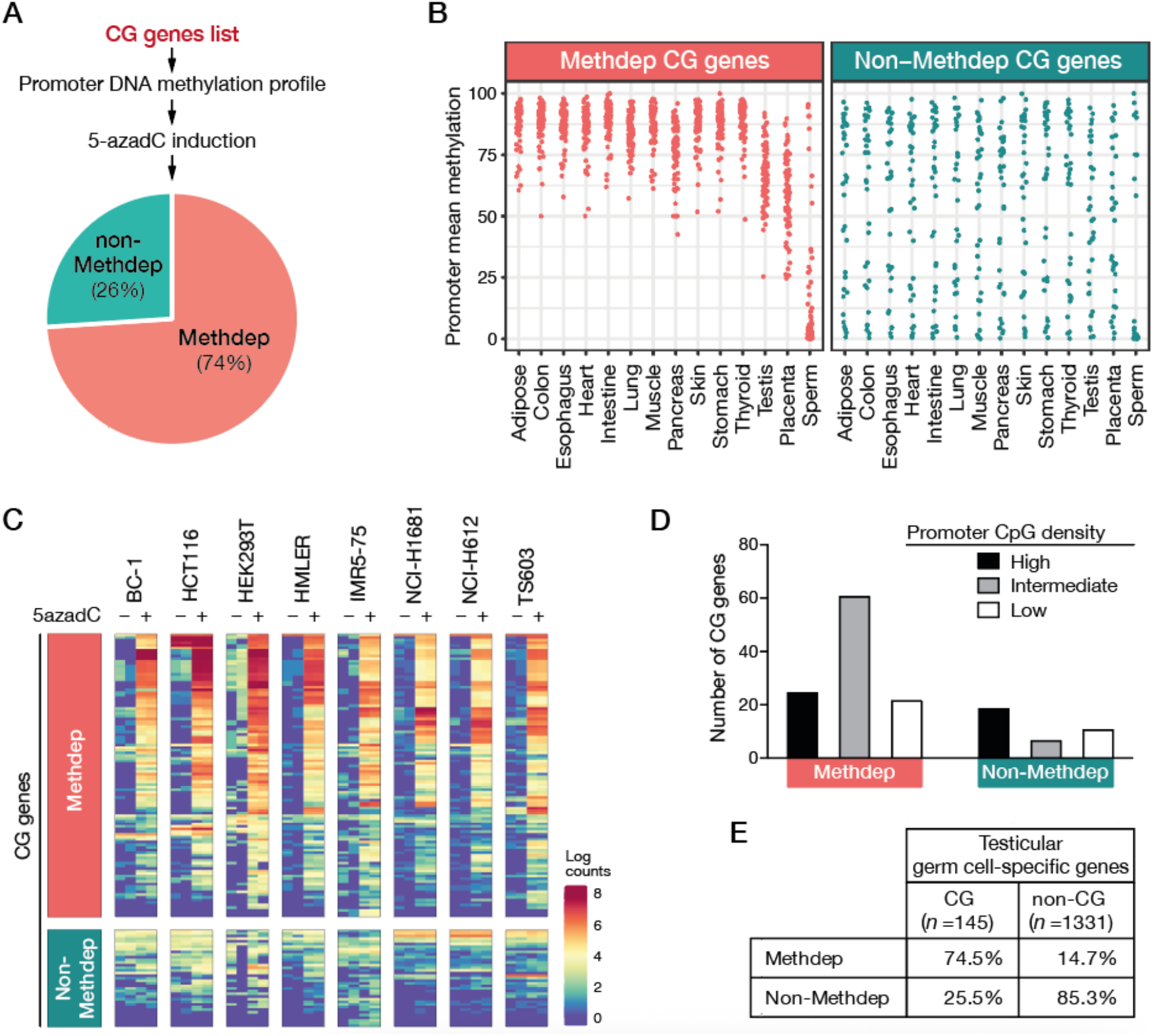
Determining DNA methylation dependency of CG genes. **(A)** Selection criteria and proportion of CG genes categorized as DNA methylation dependent (Methdep) or non-dependent (non-Methdep). **(B)** Mean DNA methylation levels of CG gene promoters among normal tissues, inferred from WGBS analyses. Each dot represents a CG gene promoter **(C)** Evaluation of the responsiveness of CG genes to induction by a DNA demethylating agent, based on RNA-Seq data on indicated cell lines exposed (+) or not (−) to 5-Aza-2′-deoxycytidine (5azadC). **(D)** CG gene promoters of Methdep and non-Methdep were classi/ied according to the CpG density (number of CpGs per 100bp) of their promoter region: high (n≥4), intermediate (2≤n<4), low (n<2). The number of CG genes in each category is represented graphically. **(E)** Proportions of Methdep genes among CG genes (1 out of the 146 could not be classi/ied) and non-CG testicular germ cell-speci/ic genes.

We then sought to determine whether CG genes display a particular propensity for methylation dependency, or whether this feature is generally shared among all testicular germ cell-specific genes, i.e., even those that do not become activated in cancer. To this end, we applied our filtering for methylation dependency to those of the initially selected testicular germ cell-specific genes that were not retained as CG genes (n=1331). The results showed that among these genes, only 14.7% qualified as Methdep, whereas 85.3% were categorized in the non-Methdep group (**Fig. 3E**). These results show therefore strong enrichment of DNA methylation dependency among CG genes (Chi-squared test, p-value = 2.33E-64), suggesting that genes using this regulatory mechanism are particularly at risk of undergoing aberrant activation in cancer.

### DNA methylation dependent (Methdep) CG genes are enriched on the X chromosome

Chromosomal assignment of the 146 CG genes revealed that 70 of them (48%) map on the X chromosome (**Fig. 4A**). This count indicates significant enrichment on the X chromosome, even after normalization to the total number of genes per chromosome, which is in line previous observations [17]. Unexpectedly, 98% (69/70) of the CG genes located on the X chromosome belong to the Methdep category. This contrasted with non-Methdep CG genes, which do not show enrichment on the X chromosome (**Fig. 4A**,**B**). Of note, a similar observation was made when comparing Methdep and non-Methdep categories for the testis-specific genes that displayed no activation in tumors and were therefore not categorized as CG genes (**Fig. 4B**). Together, these results indicate that the bias of CG genes, and more generally of testis-specific genes, for location on the X chromosome ascribes to DNA methylation dependency rather than testis-specific expression or function. This observation does not ait into the “sexual antagonism” theory to explain enrichment of testis-specific genes on the X chromosome.

**Figure 4.**
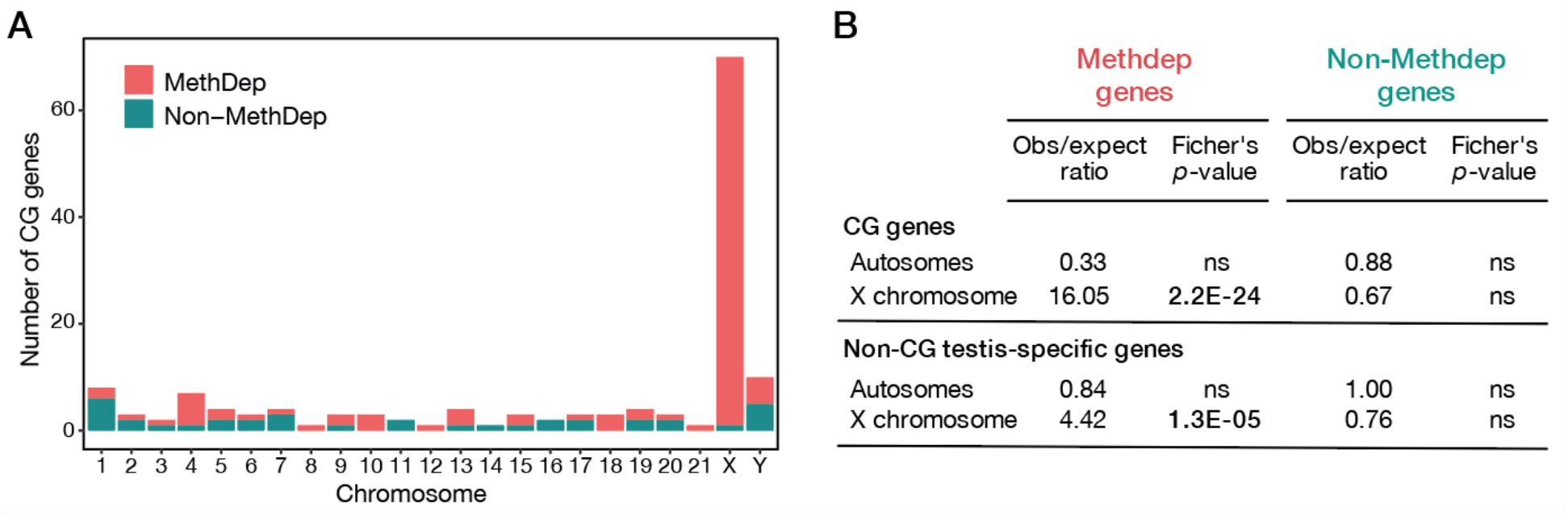
Enrichment of CG genes on the X chromosome is related to their DNA methylation dependency. **(A)** Chromosomal distribution of Methdep and non-Methdep CG genes. **(B)** Enrichment on the X chromosome in relation to DNA methylation dependency was evaluated for all testis-speci/ic genes, either CG or non-CG.

### X-linked Methdep CG genes show higher frequencies of activation in tumors

Studies have shown that CG genes vary widely in their susceptibility to become activated in tumors, with some showing activation rates that exceed 50% in certain tumors, and others never reaching more than a few percent [23]. Here, we examined if the subcategories of CG genes we defined differ in their propensity to become activated in cancer. To this end, RNA-Seq datasets of tumor cell lines (CCLE) and tissues (TCGA) of three histological types (melanoma, lung adenocarcinoma, head and neck carcinoma) were examined. We observed that CG genes of the Methdep category often display more frequent activation and higher expression levels in tumors, compared with those in the non-Methdep category (**Fig. 5A**,**B**). Highest activation frequencies and expression levels were observed for Methdep CG genes residing on the X chromosome, and especially for members of the MAGE family (**Fig. 5C**). Note that the activation frequencies of individual CG genes in all tumor types can be retrieved in the CTexploreR package.

**Figure 5.**
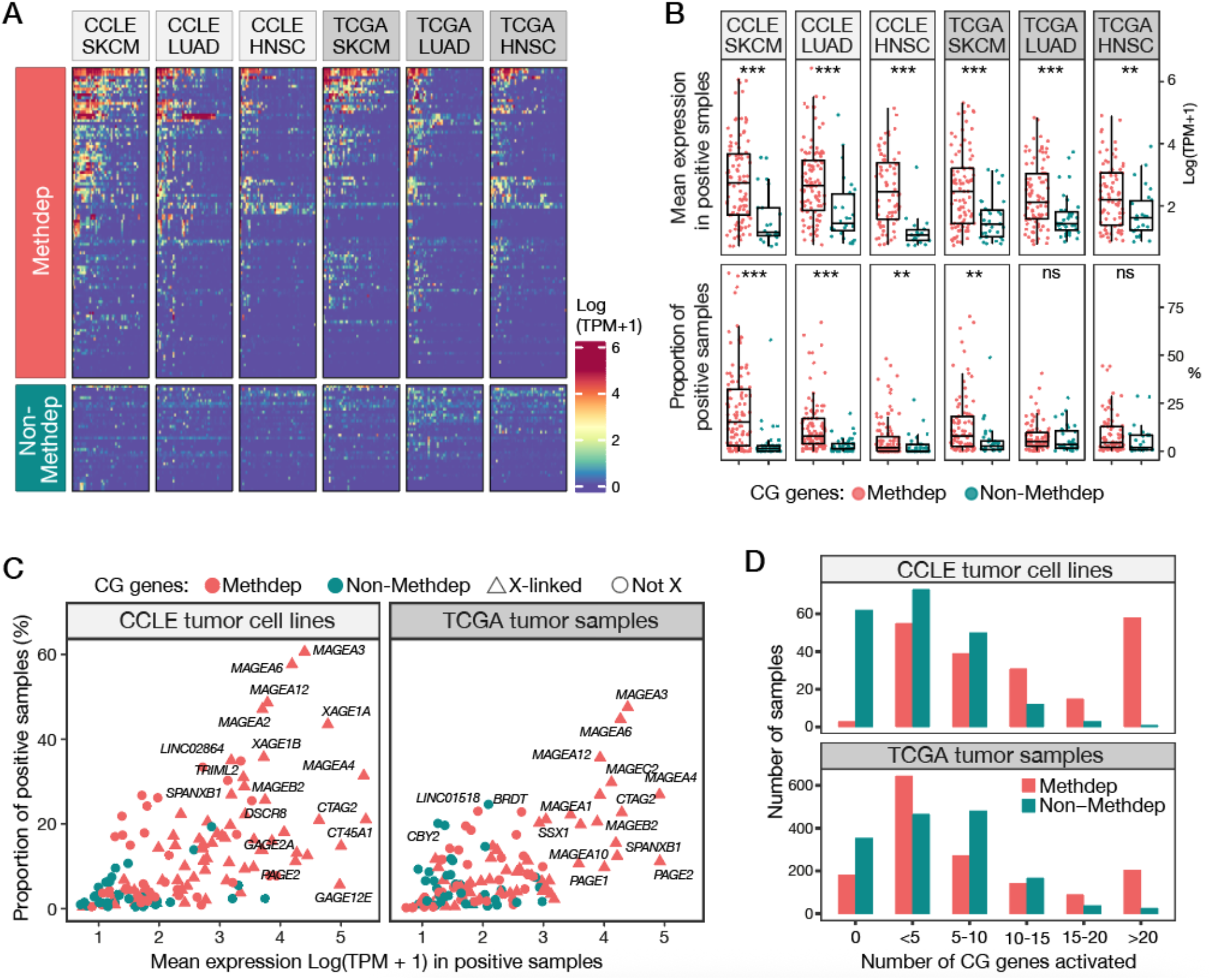
Expression of Methdep and non-Methdep CG genes in cancer cell lines (CCLE) and tumor samples (TCGA). **(A)** Heatmaps showing expression levels of CG genes in melanoma (SKCM), lung adenocarcinoma (LUAD) and head and neck carcinoma (HNSC). Fifty randomly selected cell lines or tumors are represented for each histological type. The gene order is based on Ward’s clustering. **(B)** Mean expression levels in positive samples (top panel) and activation frequencies (bottom panel) of CG genes in SKCM, LUAD and HNSC. Samples are de/ined as positive for a gene if its expression level is ≥ 1 TPM. Welch’s t-test. **(C)** As in (B) but combining the three cancer types, and identifying X-linked genes. **(D)** Comparison of co-activation of Methdep and non-Methdep CG genes in tumor cell lines and tissue samples.

CG genes are frequently found to exhibit co-activation in tumors [14,37], and this was attributed to their dependency on DNA methylation and therefore their common susceptibility to become derepressed in tumors that underwent extensive genome-wide DNA demethylation [14,39–41]. In support of this contention, we observed that Methdep CG genes show a higher tendency to become co-activated in tumor cell lines and tissues, as compared with non-Methdep CG genes which displayed more scattered patterns of activation (**Fig. 5A**,**D**). Together, these results confirm that among CG genes, those that are regulated by DNA methylation are at higher risk of becoming jointly derepressed in cancer.

### DNA methylation and chromosome location inRluence CG gene expression patterns during spermatogenesis

Previous studies have reported varying patterns of expression of CG genes during spermatogenesis, with X-linked CG genes exhibiting a marked bias towards expression in the pre-meiotic stages of germ cell differentiation [17,18,39,42]. The exact windows of expression of the different categories of CG genes have however not been systematically defined. To address this issue, we analyzed publicly available scRNA-Seq data from adult testis [43,44]. As shown in **Figure 6A**, the different categories of CG genes exhibited contrasted patterns of mRNA expression during the different stages of spermatogenesis. Most X-linked CG genes, which also belong to the Methdep category, displayed preferential expression in the pre-meiotic stages of spermatogenesis, i.e. from spermatogonia stem cells to early spermatocytes. mRNA levels of these genes decreased in late spermatocytes and became completely absent in elongated spermatids and sperm. Of note, a few X-linked Methdep CG genes exhibited a dissimilar profile of expression (*SPANXB1, SPANXD, SPANXC, DDX53, TGIF2LX*), as their mRNA became detectable from the round spermatid stage onwards up into the sperm (**Fig. 6A**). Methdep CG genes not residing on the X chromosome displayed a less restricted profile of expression, the mRNA of several of them being detected at almost all stages of spermatogenesis (**Fig. 6A**). Finally, non-Methdep CG genes (irrespective of their chromosomal location) showed varying, albeit limited, expression windows, as scRNA-Seq data identified corresponding mRNAs in germ cells representing either the pre-meiotic, meiotic, or post-meiotic stages of spermatogenesis (**Fig. 6A**). Together these results indicated that patterns of expression of CG genes during spermatogenesis is inaluenced by both dependency on DNA methylation and location on the X chromosome.

**Figure 6.**
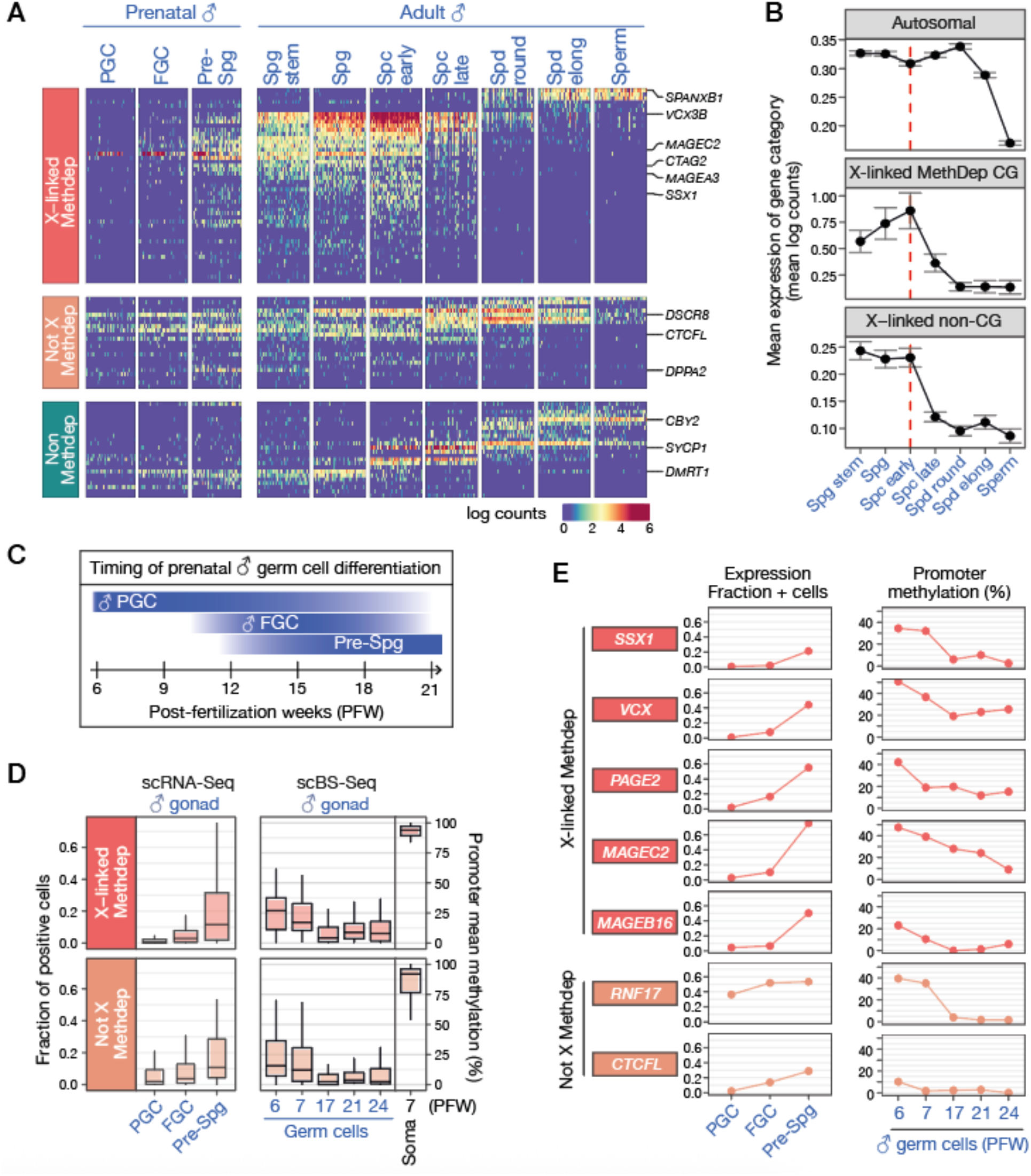
Expression and promoter DNA methylation of CG genes in the male germline. **(A)** Expression of CG genes in prenatal and adult male germ cells, based on scRNA-Seq data. Heatmaps show the data for CG genes (n=96) that were detected in ≥ 5% of any adult testis germ cell, and for 50 randomly selected cells for each germline stage (PGC: primordial germ cell, FGC: fetal germ cell, Spg: spermatognia, Spc: spermatocyte, Spd: spermatid). **(B)** mRNA expression levels (mean log counts, 95%CI) of autosomal (n = 18205), X-linked MethDep CG (n = 59) and X-linked non-CG (n = 696) genes across adult spermatogenesis. Genes undetected at pre-meiotic stages were excluded from the analysis. The red dashed line represents the onset of MSCI. **(C)** Timing of prenatal male germ cell differentiation. **(D)** Left panel: Boxplots of mean fractions of fetal germ cells where CG genes are detected (count > 0). Right panel: Boxplots of mean promoter DNA methylation levels of CG genes in male germ cells across developmental timepoints, and in male somatic cells (Soma). **(E)** Expression (fraction of positive cells) and mean promoter methylation level (%) of representative examples of CG genes in male prenatal germ cells.

It has been demonstrated that X-linked Methdep CG gene promoters maintain an unmethylated status in sperm, even though the genes are no longer expressed at this stage [16]. Post-meiotic silencing of X-linked Methdep CG genes therefore likely relies on a DNA methylation-independent mechanism. A plausible explanation is the process of meiotic sex chromosome inactivation (MSCI) taking place in spermatocytes, and leading to DNA methylation-independent downregulation of most genes residing on the X chromosome [45,46]. In support of this hypothesis, we observed that the timing of disappearance of X-linked Methdep CG mRNAs coincided with that of mRNAs originating from the bulk of genes residing on the X chromosome (**Fig. 6B**). This observation therefore suggests that location on the X-chromosome, which undergoes MSCI, accounts for the post-meiotic downregulation of X-linked Methdep CG genes.

### DNA demethylation and activation of Methdep CG genes in prenatal male germ cells

Because a number of Methdep CG genes are already expressed in spermatogonial stem cells, which correspond to the earliest germ cell stage in the adult testis, our next question was to determine whether these genes become demethylated and activated during prenatal germline development, and if so at which step. In human males, primordial germ cells (PGCs) migrate into the developing gonads around post-fertilization weeks (PFW) 4 to 6. Between PFW 8 and 10, PGC start generating germ cells that progress through successive stages of differentiation, together referred to hereafter as fetal germ cells (FGCs) (**Fig 6C**). The ultimate step of FGC differentiation results in the formation of prespermatogonia, which accumulate in the fetal gonads between PFW 12 and 20, and remain quiescent until birth [47,48]. A first wave of genome demethylation occurs in migrating PGCs, during which about 80% of DNA methylation marks are erased. This is followed by a second phase of DNA demethylation, which takes place in the gonads between PFW 7 and 11, and further reduces the proportion of methylated CpGs to 6-8% [49].

In order to assess the expression and DNA methylation status of Methdep CG genes in the different prenatal male germ cells stages, publicly available datasets from two studies that profiled the transcriptome (scRNA-Seq) and DNA methylome (scBS-Seq) at the single-cell level in prenatal human gonads (harvested between PFWs 6 and 21) were downloaded [48,50]. Upon analysis of the male gonads-derived scRNA-Seq data, we observed that the detection of mRNAs of Methdep CG genes (X and non-X) increases significantly in prespermatogonia, as compared with PGCs or FGCs (**Fig. 6A**,**D**). This observation is consistent with previous studies which identified MAGEA4, a protein encoded by an X-linked Methdep gene, as a specific marker of prespermatogonia differentiation in fetal male gonads [51]. On the other hand, examination of scBS-Seq revealed that, while the mean DNA methylation levels of Methdep CG gene promoters are already low (median = 23%) in the earliest embryonic germ cell stages (PFW 6-7), they display further decrease (median = 7%) from PFW 17 onwards (**Fig. 6C**,**D**). The latter time window follows the second genome DNA demethylation and coincides with the accumulation of prespermatogonia. Detailed examination of scRNA-seq and scBS-Seq data for individual Methdep CG genes confirmed that transcriptional upregulation in prenatal male germ cells coincides with extensive promoter demethylation (**Fig. 6E**). Together, these data suggest that, although Methdep CG gene promoters are already partially demethylated in PGCs, many of them undergo further DNA demethylation after PFW 7, and this is associated with increased mRNA expression in prespermatogonia (**Fig. 6D**). Patterns of DNA methylation and expression of individual CG genes during germ cell development can be retrieved in CTexploreR.

### Limited DNA demethylation and activation of X-Methdep CG genes in female germ cells

Germ cell development in females differs substantially from that in males, especially because meiosis is initiated during fetal development (from PFW 12), generating oocytes that remain arrested in the first meiotic prophase. Starting at puberty, subsets of oocytes in the ovaries resume maturation at each menstrual cycle, leading to the ovulation of an oocyte that further progressed up to the metaphase of meiosis 2 (MII oocyte). Completion of meiosis only occurs after fertilization of the MII oocyte. We searched to determine if the CG genes for which we detected expression in male germ cells, are also expressed in female germ cells. To this end, we used scRNA-Seq data generated from germ cells of prenatal female gonads [48], and from adult oocytes at different stages of maturation [43]. The results showed that mRNAs of 44% of the CG genes that were identified on the basis of their expression in male germ cells, were not detected in prenatal and/or adult female germ cells (compare **Fig. 6A** and **Fig. 7A**). Intriguingly, this male-biased expression was significant only for X-linked Methdep CG genes (Chi-squared test, *p*value = 1.99E-7), but not for the two other categories of CG genes (**Fig. 7B**). There were however some exceptions among X-Methdep CG genes that displayed detectable mRNA expression in female germ cells, either during prenatal (*PAGE2, PAGE5*) or adult (*MAGEB1, MAGEB2, MAGEC1, FTHL17*) stages of oocyte development (**Fig. 7A**).

**Figure 7.**
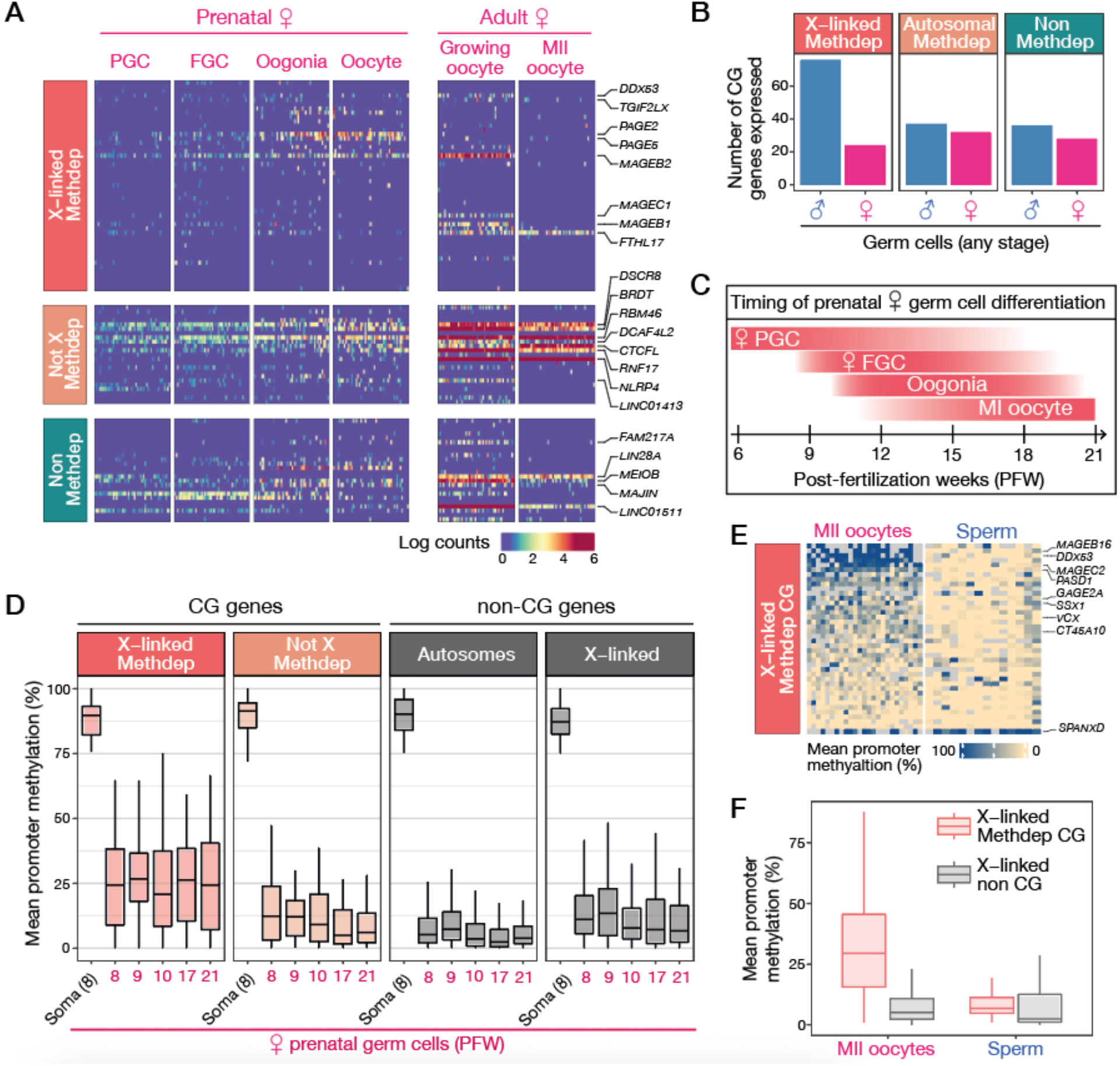
Expression and promoter DNA methylation of CG genes in the female germline. **(A)** Heatmaps showing expression of CG genes (same genes as in Fig. 6A) at different stages of prenatal and adult female germline development, based on scRNA-Seq data (50 randomly picked cells per stage are shown). **(B)** Number of CG genes expressed (count > 0 in > 10% of cells in scRNA-Seq data) in male or female germ cells of any stage. **(C)** Timing of prenatal female germ cell differentiation. **(D)** Boxplots of mean promoter DNA methylation levels of CG genes in female germ cells across developmental timepoints, and in female somatic cells (Soma). The group of non-CG genes includes 4047 automosal and 137 X-linked genes, which were selected on the basis of high DNA methylation levels (> 75%) in somatic cells. **(E)** Mean promoter DNA methylation levels of X-linked CG genes in mature MII oocytes and sperm cells (excluding Y chromosome carrying cells). **(F)** Comparison of mean promoter DNA methylation levels (%) between Methdep CG (n = 40) and non-CG (n = 780) genes residing on the X chromosome, in MII oocytes and in sperm cells.

We next explored the scBS-Seq data generated from human prenatal gonads [50], in order to establish the DNA methylation profiles of Methdep CG genes in female embryonic and fetal germ cells. It was reported that, like in males, PGCs in females undergo a first partial erasure of DNA methylation marks (down to ~20% methylated CpGs), which is followed by a second phase of DNA demethylation between PFWs 7 and 10 leaving only about 7% of CpGs methylated. Of important note, the inactivated X chromosome is already reactivated by PFW 4 in female PGCs [49]. We observed that in female gonads, X-linked Methdep CG genes exhibit a median level of promoter methylation level of 25% with no decrease over the post-fertilization weeks (**Fig. 7C, D**). This contrasted with the promoters of not-X Methdep CG genes, for which the median DNA methylation level decreased to 6% by PFW 17 (**Fig. 7D**). As a control, we also evaluated the methylation status of the bulk of somatically methylated, yet non-CG genes, residing either on autosomes or the X chromosome, and confirmed that the median level of DNA methylation within these genes decreases to 7% in prenatal oocytes (**Fig. 7D**). Together, these observations suggest that X-linked Methdep CG genes display the specific characteristic of escaping extensive promoter DNA demethylation during fetal oogenesis. Importantly, as they do not escape demethylation during male germline development (**Fig. 6D**), X-linked Methdep CG genes were expected to exhibit sex-specific DNA methylation profiles in fertilizing gametes. This was confirmed by the examination of scBS-seq datasets generated from adult sperm and MII oocytes [52], which showed that DNA methylation levels of X-linked Methdep CG gene promoters are generally higher in MII oocytes than in sperm (36.7 % vs 11,1%, Welch t-test, *p*value = 1,83E-08) (**Fig. 7E,F**). In comparison, non-CG genes residing on the X did not show similar preservation of promoter DNA methylation in MII oocytes (**Fig. 7F**).

### Sex-speciRic imprinting and expression of X-Methdep CG in early embryos

It has been shown that most DNA methylation marks carried by gametes (with the notable exception of those on parentally imprinted genes) become rapidly erased after fertilization [53]. Here, we sought to find out if DNA methylation marks on X-linked Methdep CG genes, which we found to be specifically present in oocytes, are preserved after fertilization in the embryo. If this was indeed the case, one would expect that in male embryos, which inherit only one oocyte-derived maternal X chromosome, X-linked Methdep CG genes would maintain a higher level of DNA methylation than in female embryos, which inherit two X chromosomes, one of maternal and the other of paternal origin. To address this issue we analyzed scBS-Seq datasets generated from mature human gametes and cells isolated from pre-implantation embryos at different times post-fertilization [52]. Interestingly, we found that X-linked Methdep CG genes exhibit significantly higher promoter DNA methylation levels in male versus female embryos, from the zygote to the blastocyst stage (**Fig. 8A**), thereby supporting preservation of oocyte-specific DNA methylation on these genes during early embryonic development. After implantation, X-linked Methdep CG genes showed extensive de novo DNA methylation in both male and female embryos, although they maintained a promoter methylation level that was slightly higher in male embryos (**Fig. 8A**). Detailed analysis of scBS-Seq data for individual X-linked Methdep CG genes confirmed male-biased DNA methylation in pre-implantation embryos, and also revealed allele-specific DNA methylation profiles in XX female embryos, most likely realecting the presence of an unmethylated allele of paternal origin and a methylated allele of maternal origin (**Fig. 8B**).

**Figure 8.**
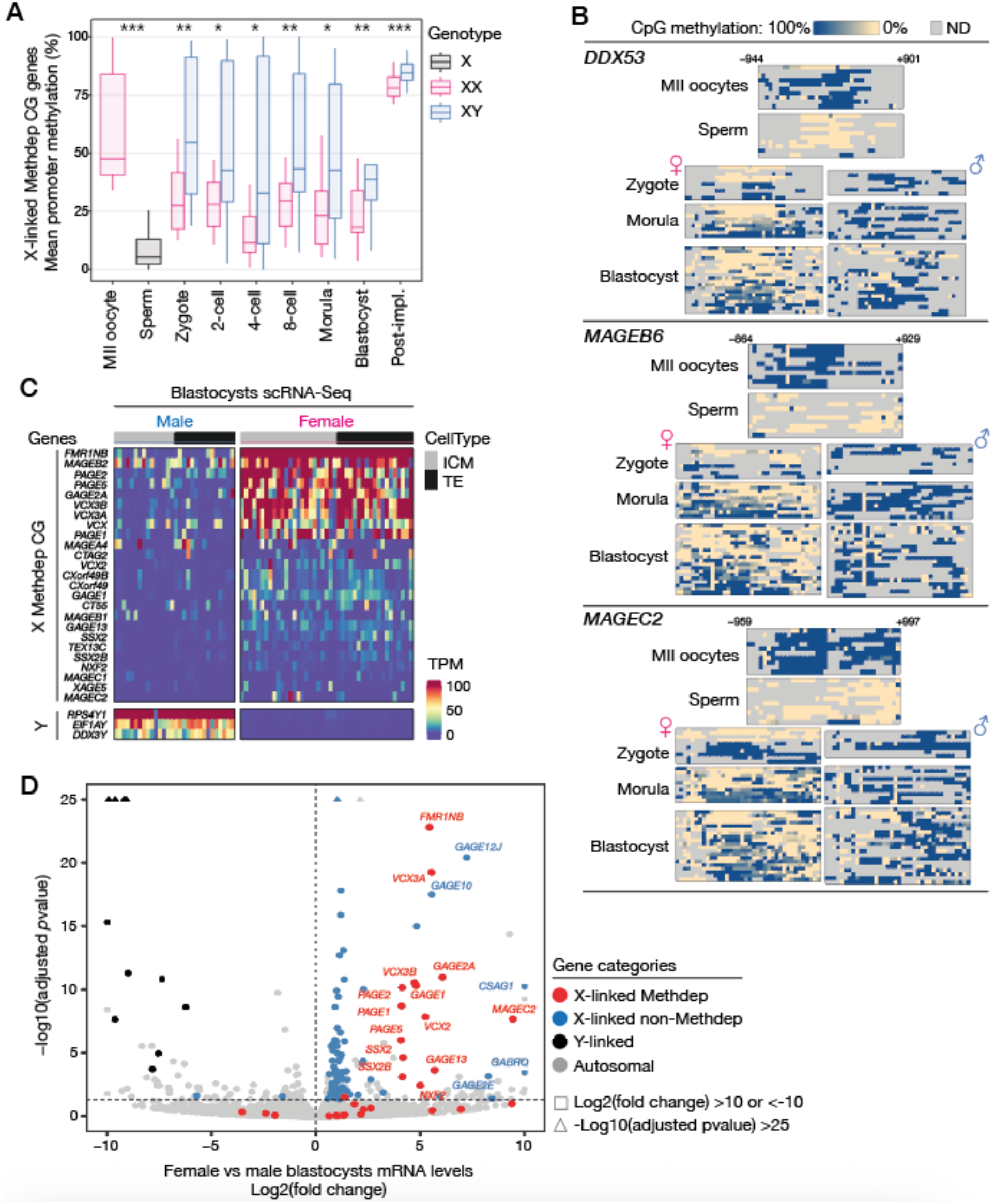
Transient maternal imprinting and sex-speciOic expression of X-linked Methdep CG genes in preimplantation embryos. **(A)** scBS-Seq data of germ cells and early embryos (male or female) were analyzed to evaluate mean promoter DNA methylation levels of X-linked Methdep CG genes (only those initially showing >33% methylation in MII oocytes). Y-carrying sperm cells were excluded from the analysis. Welch’s t-test. **(B)** Detailed scBS-Seq results of representative Methdep X-linked CG genes in male and female germ cells and preimplantation embryos. **(C)** Expression (TPM) of X-linked Methdep CG genes in individual cells of the inner cell mass (ICM) or trophectoderm (TE) of male and female blastocysts. The heatmap shows only genes that were detected in > 20% of cells of any type. Y-linked genes are shown as controls. **(D)** Volcano plot representing the result of a global differential expression analysis comparing female and male blastocyst cells.

We next searched to determine if the sex-specific DNA methylation patterns we observed are correlated with differential expression of X-linked Methdep CG genes in male and female embryos. It was to be expected, indeed, that these genes would remain silent in XY male embryos, which inherit only a maternal X chromosome on which they are methylated. In female embryos, on the other hand, the presence of an X of paternal origin should allow their expression. Analysis of scRNA-Seq data generated from human blastocysts of both sexes [52], demonstrated that about 40% of X-linked Methdep CG genes show preferential expression in female embryos, where higher levels of their mRNA were detected in cells from both the inner cell mass and the trophectoderm (**Fig. 8C**). Furthermore, genome-wide differential expression analysis showed that X-linked Methdep CG genes stand out from all other genes by their preferential expression level in female embryos (**Fig. 8D**, Supplemental **Fig. S2**). Of note, it is unlikely that this expression bias is solely attributable to double gene dosage in XX embryos, as the level of female-biased expression (fold change) of X-linked Methdep CG genes was markedly higher than that of other genes on the X chromosome (**Fig. 8D**). A few X-linked genes that had not been categorized as CG genes (eg. *GAGE12J, GAGE10, CSAG1*) nevertheless showed high female-preferential expression. These genes actually belong to well-described CG gene families (*GAGE, CSAG*), but did not pass our highly stringent criteria of inclusion into the CG group of genes. Together, our analyses uncovered an intriguing new feature of human X-linked Methdep CG genes, by demonstrating that some of them carry transient sex-specific imprints, and display therefore female-biased expression in early embryos. Of important note, the analysis of scRNA-Seq data generated from murine embryos [54] revealed that this sex-specific expression does not occur in the mouse (**Fig. S3**). This can be explained by the lack of conservation of CG genes in the mouse (**Fig. S3**), and by the existence of a process of paternal X chromosome inactivation occurring in early female mouse embryos [55].

## Discussion

We created CTexploreR, a R/Bioconductor package, to generate an updated database describing CG genes. We used omics data to define a list of 146 CG genes, based on stringent criteria to select genes that become activated in cancerous lesions, but otherwise display highly restricted expression in testicular germ cells. These genes and their main characteristics are listed in a table that constitutes the core of CTexploreR. Additionally, specific functions have been created to facilitate access to all the data presented in this study, allowing users to quickly visualize (or extract) mRNA expression and DNA methylation profiles of any CG gene in normal or tumoral tissues and cells. Of note, these functions can be applied to the analysis of any human gene, extending the utility of CTexploreR to users wishing to test their favorite gene in any of the transcriptomic and methylomic datasets.

Our main objective with CTexploreR was to analyze CG genes in a systematic way, in order to investigate the existence of gene subgroups differing on the expression pattern, epigenetic regulation, and chromosomal location. A well-documented characteristic of CG genes is their reliance on DNA methylation as a primary mechanism of repression in somatic tissues [17,53]. There is evidence, however, suggesting that this may not apply to all CG genes [17,56]. The present study supports the view that DNA methylation is indeed involved in the regulation of a majority of CG genes (74% Methdep CG genes). Yet, a fraction of CG genes do not seem to be directly controlled by DNA methylation (26% non-Methdep CG genes). Contrary to Methdep CG genes, which were often co-activated in tumors, non-Methdep CG genes exhibited scattered patterns of expression among tumor samples, suggesting that their activation in tumors relies on gene-specific regulatory mechanisms rather than a shared process of transcriptional de-repression.

Among the list of 146 CG genes, 48% mapped on the X chromosome. This enrichment is consistent with the previously reported observation that genes with testis-specific expression are generally over-represented on the X chromosome in mammals [57,58]. The explanation put forward relies on the theory of sexual antagonism, which posits that genes beneficial to males and detrimental to females accumulate preferentially on the X chromosome, because X hemizygosity in males allows recessive mutations to aix more efficiently on this chromosome [10,59]. Surprisingly, we observed that enrichment on the X chromosome concerned only the testis-specific genes that display DNA methylation dependency. Genes owing their testis-specific expression to mechanisms other than DNA methylation showed unbiased chromosomal distribution. It seems therefore that the accumulation of testis-specific genes on the X chromosome is not merely associated with male-beneficial functions, but also with their DNA methylation-based mode of transcriptional regulation. The reason for this is unclear, but may be linked with the evolutionary origin of X-linked CG genes, many of which present as multigene families that were recently acquired through palindromic duplications [11,60–62]. Compared to autosomes, the X chromosome shows an enrichment in palindromic duplications [63,64]. This may be explained by an increased propensity of the X chromosome to undergo rearrangements during male meiosis, resulting from the absence of a fully homologous pairing partner during meiotic recombination [63,65]. Interestingly, it has been shown that duplicated DNA segments often show high levels of CpG methylation [66,67], thereby suggesting that X-linked CG genes may have acquired these marks at the time of their formation. It is expected that with this methylated state, X-linked CG genes spontaneously adopted specific expression in germ cells, due to the process of genome DNA demethylation that occurs in the germline during development [49,68,69]. In brief, we propose that CG genes on the X chromosome have multiplied as a result of duplication processes that concur with DNA methylation, and hence adopted “by default” a testis-specific expression pattern dictated by global epigenetic aluctuations in germ cells.

X-linked Methdep CG genes were for the most part silenced from the spermatocyte stage onwards. This coincides with the well described process of meiotic sex chromosome inactivation (MSCI), which leads to DNA methylation-independent downregulation of most genes residing on the X chromosome during meiosis 1 [45,46]. Location of Methdep CG genes on the X chromosome appears therefore as a necessary feature to limit their expression during the pre-meiosis and meiosis 1 stages of spermatogenesis. This may be another contributing factor to the enrichment of these genes on the X chromosome.

Contrasting with X-linked CG genes, CG genes residing on autosomes are mostly single-copy, and do not exhibit the evolutionary novelty of those located on the X chromosome [11,70]. Our analysis indicated that half of non-X CG genes do not seem to depend on DNA methylation for their tissue-specific expression. These Non-Methdep CG genes had variable activation profiles in the adult testis, where they showed narrow expression windows in defined stages of spermatogenesis. Such particular expression profiles are likely imposed by specific sets of transcription factors that become activated sequentially before and during male meiosis, as described previously [71].

Although previous studies have reported the expression of several CG genes during oogenesis [48], our study shows that about half of CG genes are exclusively expressed during male gametogenesis. This was particularly true for the subgroup of X-linked Methdep CG genes, and was consistent with our observation that, whereas the promoters of these genes become fully demethylated in male fetal germ cells, they resist DNA demethylation during female germ cell development. Of important note, persistent DNA methylation on these gene promoters in female germ cells cannot be ascribed to X chromosome inactivation, since the inactive X chromosome is reactivated at a much earlier stage of embryogenesis [49]. Thus, X-linked Methdep CG genes belong to the few genomic sequences that escape the process of genome-wide DNA demethylation in developing female germ cells [49,50,69]. The mechanisms that allow X-linked Methdep CG genes to resist DNA demethylation during female germline development remain to be discovered. Importantly, because X-linked Methdep CG genes escape DNA methylation reprogramming during female oogenesis, they may contribute to the transmission of an epigenetic inheritance from the mother to her offspring.

The most intriguing finding of our study is that X-linked Methdep CG genes display female-biased expression in the early embryo. This results from the differential aluctuation of their promoter DNA methylation levels during male and female germline development, and the preservation of maternally inherited DNA methylation marks during early embryo development (**Fig. 9**). Thus, X-linked Methdep CG genes exhibit transient maternal imprinting, which later becomes obliterated when the paternal allele also becomes de novo DNA methylated at around the time of implantation. Transient imprints are not uncommon in early embryos [72–76], but have moderate impact on mRNA levels for genes residing on autosomes, since epiallelic forms of both parents are present. The consequence of transient maternal imprinting is radically different when it concerns genes located on the X chromosome, because XY male embryos inherit only the maternal epiallele. This is indeed what we observed for X-linked Methdep CG genes, which displayed strong female-biased expression in early stage embryos (**Fig. 9**). Genes displaying sex-differential expression in early embryos are a source of great interest [75,77,78], because they challenge the notion that sexual differentiation is initiated only after the 6^th^ week of gestation, when the master regulator of male gonad differentiation (SRY) starts to be expressed [79,80]. Our transcriptomic analyses in early embryos indicated that X-linked Methdep CG genes show the highest female-biased expression of all genes, suggesting that they may have an important impact in preimplantation sexual dimorphism. Even more remarkable is the fact that X-linked Methdep CG genes exhibit male-biased expression in germ cells, and instead female-biased expression in early embryos (**Fig. 9**).

**Figure 9.**
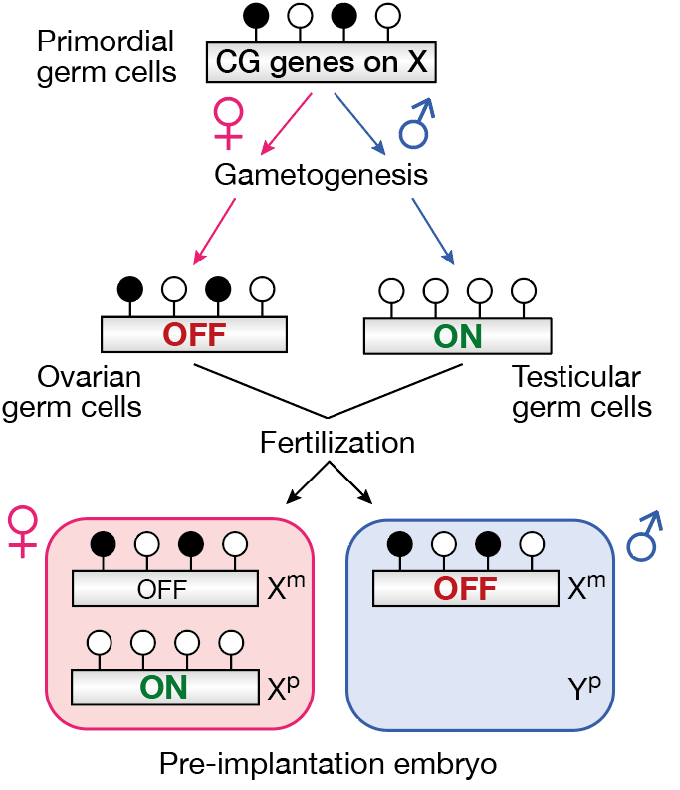
Maternal imprinting of X-linked Methdep CG genes during gametogenesis results in female-speciOic expression in pre-implantation embryos. The parental origin of the chromosomes that are inherited in pre-implantation embryos is indicated: maternal (m), paternal (p).

It is difficult to predict how X-linked Methdep CG genes may impact early embryo biology, because their cellular functions were exclusively investigated in spermatogenic and cancer cells, and remain for the most part ill-defined. Several members the *GAGE* gene family (*GAGE1, GAGE2A, GAGE12J*) were among the most differentially expressed genes when we compared male and female embryos. A study in human cancer cells showed that GAGE proteins, and in particular GAGE12 variants, recruit histone deacetylases on the chromatin, and induce local decompaction to facilitate DNA repair [81]. As a result, tumor cells that overexpress *GAGE* genes were found to be more resistant to irradiation- or chemotherapy-induced DNA damage [81,82]. Genes belonging to the *MAGE* family (*MAGEB2*,*B6* and *MAGEC2*) also displayed substantial female-biased expression in early embryos. Previous experiments in tumor cell lines revealed that MAGE proteins modulate the activity and/or stability of P53 and AMPKα1, two master sensors governing cellular stress response and metabolic adaptation, respectively [83–85]. Consistently, gene depletion experiments in the mouse revealed that Mage proteins exert a protective role in spermatogenic cells of animals that are exposed to a genotoxic stress or to long term starvation [86]. If the genes described here above exert similar functions during early embryogenesis, one might expect that female embryos will be better protected than male embryos against environmental stress. These CG genes could therefore have an impact on the sex ratio of developing embryos when pregnancy occurs in unfavorable conditions. Interestingly, it has been shown that mothers enduring famine show a sex-ratio that shifts towards more female births, and the same trend was observed in gestating animals that were experimentally undernourished [87,88].

In summary, our work revealed that, on the basis of DNA methylation dependency and chromosomal location, CG genes can be divided into three subgroups, which differ in their pattern of expression in tumors, germ cells, and early embryos. We also found that the well-known over-representation of testis-specific genes on the X chromosome applies specifically to genes that rely on DNA methylation for silencing in somatic tissues, which brings in the need to reconsider the evolutionary basis of this enrichment. Finally, we show that, due to differential DNA methylation reprogramming during male and female gametogenesis, a set of CG genes display sex-biased expression in early embryos. Therefore, investigations aiming at understanding the function of CG genes, and hence their potential contribution to tumor development, should take into consideration not only their role in spermatogenesis, but also their contribution to sex-specific embryonic development.

## Material and Methods

### Selection of CG genes

#### 1. Selection of testis-speciHic and testis-preferential genes using bulk RNAseq data from normal tissues

RNAseq expression data from normal tissues were obtained from the GTEx portal [84] (GTEx_Analysis_2017-06-05_v8_RNASeQCv1.1.9_gene_median_tpm.gct.gz) and used to identify testis-specific and testis-preferential genes. Testis-specific genes were defined as genes expressed in testis (TPM ≥ 1) but not in somatic tissues (TPM < 0.5 in all somatic tissues), and with an expression value in testis at least 10 times higher than in any somatic tissue. Testis-preferential genes were defined as genes expressed in testis (≥ 1 TPM), silent in at least 75% of somatic tissues (TPM < 0.5), and for which a certain level of expression (always at least 10x lower than the level detected in the testis) was tolerated in a minority (less than 25%) of somatic tissues. Additionally, genes that were initially undetectable in the GTEx database (TPM < 1 in all tissues) were tested in another dataset of normal tissues. We used RNAseq fastq files of 18 normal tissues that were downloaded from ENCODE [89] (accession numbers are listed in supplemental **Table S3**) that were reprocessed (as described in the RNAseq processing section) to include or not multi-mapped reads in the counting step. Genes that became detectable in testis (TPM ≥ 1) when multi-mapped reads were counted (at least 5x more than when multi-mapped reads were discarded) but remained low in somatic tissues (expression at least 10x lower than in testis) were classified as testis-specific if their TPM value was <1 in all somatic tissues or as testis-preferential otherwise.

#### 2. Filtering of germline-speciHic genes using scRNA-Seq data from normal tissues

The Single Cell Type Atlas classification (https://www.proteinatlas.org/download/proteinatlas.tsv.zip) from the Human Protein Atlas [31] was used to exclude from our initial selection of testis-specific genes the ones that were alagged as specific of any somatic cell type, including somatic cells originating from the testicular tissue. Genes that were excluded because they had low expression in some somatic cell types (at least 10x lower than the level detected in any germ cell type) were re-classified as testis-preferential genes.

#### 3. Selection of genes activated in tumors

RNA-Seq data from The Cancer Genome Atlas (TCGA) [85] was downloaded using TCGAbiolinks v2.25.3 [90] and used to test the activation of selected genes in 4141 tumor samples corresponding to seven different tumor types (SKCM, LUAD, LUSC, COAD, ESCA, BRCA, and HNSC). Genes detected (TPM > 1) in at least 1% of tumors and displaying an expression value higher than 5 TPM in at least one tumor sample were selected. To make sure that any expression really originates from a cancer cell rather than the tumor microenvironment, we also applied the same activation criteria using RNA-Seq data from the Cancer Cell Line Encyclopedia (CCLE) [86] (data and metadata were downloaded from https://ndownloader.aigshare.com/ailes/34989922 and https://ndownloader.aigshare.com/ailes/35020903). 1229 cancer cell lines originating from 20 different tissue types were tested. The 1% activation threshold we choose was justified by the fact that a number of well-known CG genes were detected in only 1-3% of tumor cell lines.

#### 4. Additional Hiltering of leaky genes

TCGA and CCLE data were also used to filter out leaky genes that passed through our initial selection criteria of testis-specific genes. We reasoned that a gene not completely silent (TPM < 0.5) in at least 20% of the tumor samples and cancer cell lines is at risk to be constitutively expressed rather than induced by a tumor-associated activation process. These genes were removed. Similarly, we made use of the normal peritumoral samples available in TCGA data to remove from our selected genes those that were already detected in a significant fraction of these cells (TPM > 0.5 in more than 25% of normal peritumoral tissues).

#### 5. Manual curation

All selected genes were visualized using IGV (Integrative Genome Viewer) [91] and a bam alignment file from testis. Unexpectedly we observed that for some genes, the reads were not properly aligned on exons but were instead spread across a wide genomic region spanning the genes, likely realecting a poorly defined transcription. These genes were removed from our selection.

#### 6. CG genes on the Y chromosome

CG genes located on the Y chromosome (n=10), which are present only in male individuals, were ignored in subsequent analyses evaluating expression and DNA methylation patterns in tumors, germ cells and pre-natal stages of both sexes.

### Methylation dependency analyses

WGBS data of 13 normal tissues were downloaded as bed files from ENCODE [89] (accession numbers are listed in supplemental **Table S3**). Additionally, a sperm WGBS fastq file was downloaded from SRA (accession SRR15427118) and processed using Trim Galore v0.5.0 to trim adapters and remove low quality reads. Methylation calling was done with Bismark v0.20.0 [92]. Methylation values corresponding to the same CpG sequenced in forward and reverse sense were averaged. Promoter methylation levels represent the mean methylation values of all CpGs located 1 kb upstream, and 200 pb downstream each transcription start site (TSS). For genes displaying multiple TSS, we used the TSS of the canonical transcript retrieved from the Ensembl database using biomaRt package v2.54.0 [93].

We also used RNA-Seq fastq files of 8 cell lines treated or not with the DNA methylation inhibitor 5-Aza-2′-deoxycytidine (5-AzadC), which were downloaded from SRA (accession numbers are listed in supplemental **Table S3**). The files were processed as described in the RNA-Seq processing section, allowing the counting of multi-mapped reads. Differential expression analyses were performed in each cell line using DESeq2 (v1.46.0) [94], to identify genes significantly up-regulated by 5-AzadC.

Two criteria were used to classify genes as methylation dependent: first, they had to be significantly induced by the demethylating agent in at least one of the 8 cell lines (logFC > 2 and p-adjusted value < 0.1). Secondly, their promoter had to be highly methylated in normal somatic tissues (mean methylation in all somatic tissues > 50%). When WGBS data was missing for a gene, only the second criteria was used for the classification.

CTexploreR provides a measurement of the density of CpGs within each promoter (−1000bp to +200bp), expressed as number of CpGs /promoter length x 100 (CpG_density). On this basis genes were classified into three categories of promoter CpG densities (CpG_promoter): “low” (CpG_density < 2), “intermediate” (2 ≤ CpG_density < 4), and “high” (CpG_density ≥ 4).

### RNA-Seq processing

The quality of fastq files was assessed using FastQC (v0.11.8) [95] and Trimmomatics (v0.38) [96] was used remove low quality reads and adapters. Reads were aligned on grch38 genome using hisat2 (v2.1.0) [97] and gene expression levels were evaluated using featureCounts from Subread (v2.0.3) [98] and Homo_sapiens.GRCh38.105.chr.gtf.gz gtf file. Both software were launched with default settings to discard multi-mapped reads. To allow multi-mapping, the -k parameter of hisat2 was set to 20 (to increase the number of primary alignments reported) and featureCounts was run with -M parameter (to count reads aligned to multiple locations).

### scRNA-Seq datasets

scRNA-Seq data from human adult testis and oocytes were downloaded from GEO (accessions GSE112013 and GSE154762) and fetal gonads scRNAseq data was downloaded from (https://cellxgene.cziscience.com/collections/661a402a-2a5a-4c71-9b05-b346c57bc451Data).

Raw counts and annotations provided by the authors were used and stored as a SingleCellExperiment object [99]. Count values were normalized and log-transformed using the logNormCounts function of the scuttle package [100]. Two scRNA-Seq datasets from blastocysts were used to compare male and female embryos (see the differential expression analysis section below). For the first one (from [52]), as raw counts were not available, fastq files were downloaded from SRA (accession SRP074598) and reprocessed as described in the RNA-Seq processing section. Cells with less than 7E+06 reads and less than 7500 detected genes were considered as outliers and were removed. Raw counts and metadata of the second dataset (from [101]), were downloaded from ArrayExpress (E-MTAB-3929).

### scBS-Seq datasets

DNA methylation data from fetal germ cells were downloaded as bed files (hg19 coordinates) from GEO (accession GSE107714). Oocytes, sperm and early embryos methylation files (hg19 coordinates) were downloaded from GEO (accession GSE81233). The mean methylation of each promoter was calculated at each stage or timepoint using methylation values of all CpG located 1 kb upstream, and 500 pb downstream each TSS (whose coordinates were converted to hg19 assembly using liftOver [102]).

### Differential expression analyses in male and female embryos

Blastocysts scRNA-Seq raw counts from [52] were summed by embryo ID and by cell-type (ICM or TE derived cells) using the summarizeAssayByGroup function of the scuttle package [100]. This led to 5 ICM-derived pseudo-bulk samples (2 males and 3 females) and 5 TE-derived pseudo-bulk samples (2 males and 3 females). DESeq2 (v1.46.0) [94] was used to perform a differential expression analysis comparing male and female samples, using the design ~ sex + cell-type. Similarly, the second blastocyst dataset (from [101]) was used to produce the volcano plot represented in supplemental **Fig S2**. Counts from embryos between E4 and E7 were aggregated by embryo ID, by day and by lineage (epiblast, primitive endoderm, trophectoderm or not applicable). Pseudo-bulk samples with less than 3 cells were removed. DESeq2 (v1.46.0) [94] was run using the design ~ sex + lineage + day.

### CTexploreR package

CTexploreR (www.bioconductor.org/packages/release/bioc/html/CTexploreR.html) has been developed as an exploratory visualization tool, allowing users to generate quick representations of CG genes expression and methylation in a series of normal and cancer samples [98]. It was developed together with a companion package, CTdata [103], that stores all the datasets used by CTexploreR. Both CTexploreR and CTdata are reviewed R/Bioconductor packages.

### Statistical analyses and graphical representations

Data analyses were done using R programming language (v 4.4.0). Figures were generated with ggplot2 v3.5.1 [104], UpSetR [105] and ComplexHeatmap v2.22.0 [106]. The entire code used to generate figures in this article is available on the GitHub repository (https://github.com/UCLouvain-CBIO/2023-CTexploreR). In some figures, statistical significance is represented by asterisks (*/**/*** indicates *p* value < 0.05/0.01/0.001, respectively, ns indicates non-significant).

## Supporting information

**Figure S1. Enabling multimapping in RNA-Seq data analysis reveals members of multigene families that were otherwise invisible**.

**Figure S2. Volcano plot of a differential expression analysis comparing female versus male preimplantation embryonic cells**.

**Figure S3. Little conservation and lack of sex-biased expression of X-Methdep CG genes in the mouse**.

**Table S1: List of 146 CG genes**

**Table S2: List of 134 CG-Preferential genes**

**Table S3. List of datasets used in CTexploreR analyses**

