## Supplemental Data 1 for "A survey of human cancer-germline genes : linking X chromosome localization, DNA methylation and sex-biased expression in early embryos"

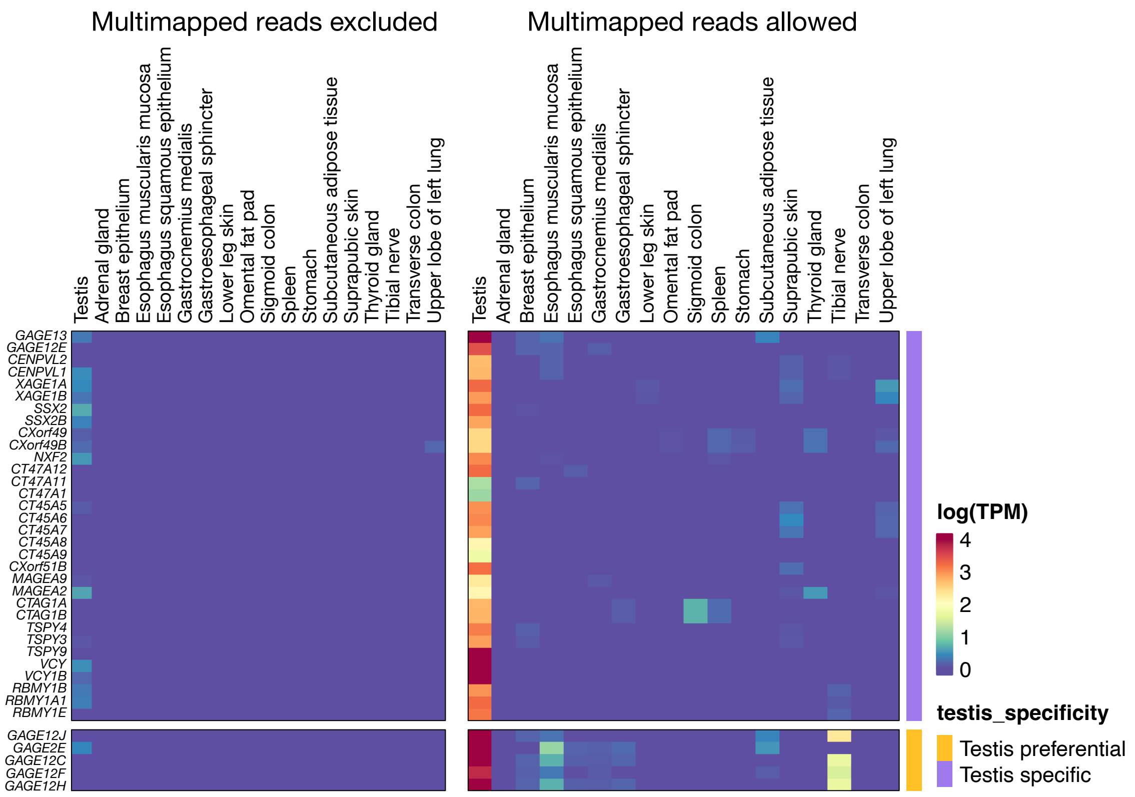


**Figure S1. Enabling multimapping in RNA-Seq data analysis reveals members of multigene families that were otherwise invisible.** CG genes that were missing in the GTEx analysis, were re-analyzed using raw RNA-Seq data of normal human tissues. Heatmaps compare results depending on whether multimapping was excluded (left) or allowed (right) during counting of RNA reads. “Testis-specific“ genes were taken into account for classification into the group of CG genes (Table S1), whereas “Testis-preferential“ were considered for inclusion into the “CG-preferential“ group of genes (Table S2).

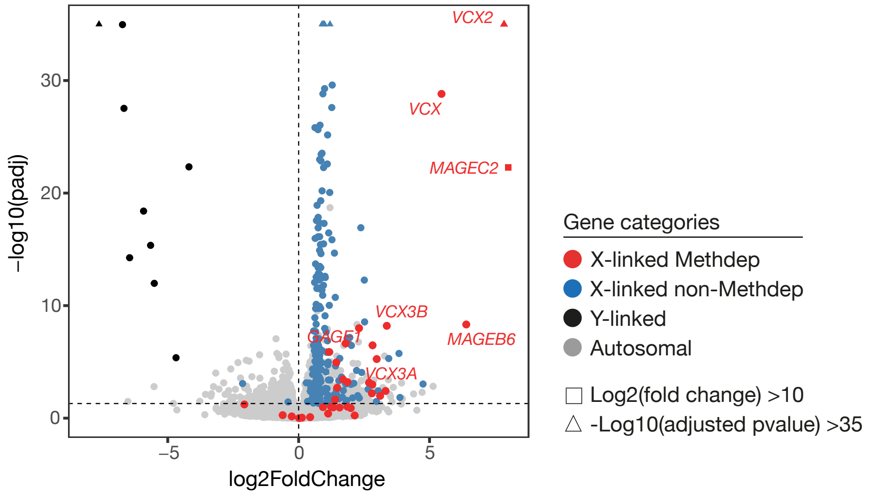


**Figure S2. Volcano plot of a differential expression analysis comparing female versus male preimplantation embryonic cells.** The analysis was based on scRNA-Seq data generated by Petropoulos and colleagues (Petropoulos et al., 2016, *Cell* 165, 1–15). The results confirm those presented in Fig. 8D, which derived from a scRNA-Seq study performed by another group (Zhu et al., 2017, Nat Genet 50, 12-19).

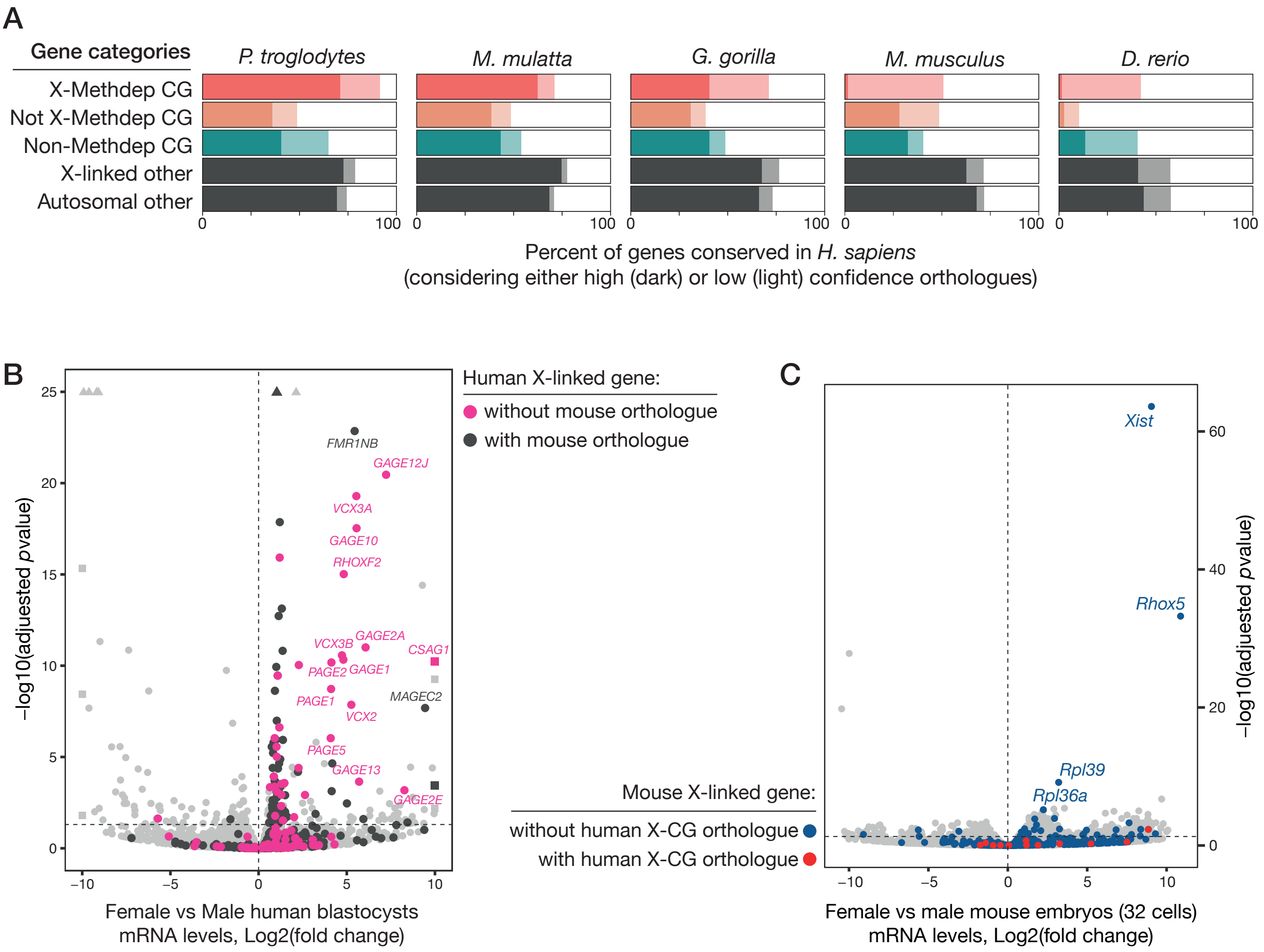


**Figure S3. Little conservation and lack of sex-biased expression of X-Methdep CG genes in the mouse. (A)** Ensembl datasets were analyze with BiomaRt to evaluate the presence of orthologues of the different categories of human CG genes in the indicated species. Conservation was compared to the bulk of X-linked or autosomal human genes, and is expressed as the percentage of human genes in the indicated category that have at least one orthologous gene in the species, using either the high or low confidence Ensembl setting. **(B)** Representation of the volcano plot of Fig. 8D highlighting human X-linked genes with or without a mouse orthologue. The figure shows that most sex-biased X-Methdep CG genes do not have a mouse orthologue. **(C)** Volcano plot representing the results of a global differential expression analysis comparing female and male mouse embryos (32-cells stage, Borensztein et al. 2017). Mouse X-linked genes that are orthologous to a human X-CG gene are highlighted in red, those that are not are in blue. Light grey dots correspond to all non-X mouse genes. The female-biased expression of *Xist* reflects the inactivation of the paternal X chromosome that occurs in early female embryos in mice (but not in humans).

**Table S1: List of 146 CG genes**

| | ensembl_gene_id | external_gene_na | CT_gene_type | testis_specificity | regulated_by_methylation | chr |
|---|---|---|---|---|---|---|
| 1 | ENSG00000234593 | KAZN-AS1 | CT_gene | testis_specific | TRUE | 1 |
| 2 | ENSG00000131914 | LIN28A | CT_gene | testis_specific | FALSE | 1 |
| 3 | ENSG00000142698 | C1orf94 | CT_gene | testis_specific | FALSE | 1 |
| 4 | ENSG00000237853 | NFIA-AS1 | CT_gene | testis_specific | FALSE | 1 |
| 5 | ENSG00000137948 | BRDT | CT_gene | testis_specific | TRUE | 1 |
| 6 | ENSG00000198765 | SYCP1 | CT_gene | testis_specific | FALSE | 1 |
| 7 | ENSG00000237463 | LRRC52-AS1 | CT_gene | testis_specific | FALSE | 1 |
| 8 | ENSG00000215817 | ZC3H11B | CT_gene | testis_specific | #N/A | 1 |
| 9 | ENSG00000162843 | WDR64 | CT_gene | testis_specific | FALSE | 1 |
| 10 | ENSG00000231532 | LINC01249 | CT_gene | testis_specific | FALSE | 2 |
| 11 | ENSG00000226856 | THORLNC | CT_gene | testis_specific | FALSE | 2 |
| 12 | ENSG00000196604 | POTEF | CT_gene | testis_specific | TRUE | 2 |
| 13 | ENSG00000163530 | DPPA2 | CT_gene | testis_specific | TRUE | 3 |
| 14 | ENSG00000242512 | LINC01206 | CT_gene | testis_specific | FALSE | 3 |
| 15 | ENSG00000251350 | LINC02475 | CT_gene | testis_specific | TRUE | 4 |
| 16 | ENSG00000170516 | COX7B2 | CT_gene | testis_specific | TRUE | 4 |
| 17 | ENSG00000250102 | LINC02377 | CT_gene | testis_specific | TRUE | 4 |
| 18 | ENSG00000250033 | SLC7A11-AS1 | CT_gene | testis_specific | TRUE | 4 |
| 19 | ENSG00000151962 | RBM46 | CT_gene | testis_specific | TRUE | 4 |
| 20 | ENSG00000231171 | LINC01098 | CT_gene | testis_specific | FALSE | 4 |
| 21 | ENSG00000179046 | TRIML2 | CT_gene | testis_specific | TRUE | 4 |
| 22 | ENSG00000250584 | LINC01511 | CT_gene | testis_specific | FALSE | 5 |
| 23 | ENSG00000251629 | LINC02241 | CT_gene | testis_specific | TRUE | 5 |
| 24 | ENSG00000164256 | PRDM9 | CT_gene | testis_specific | TRUE | 5 |
| 25 | ENSG00000251273 | LINC02228 | CT_gene | testis_specific | FALSE | 5 |
| 26 | ENSG00000145975 | FAM217A | CT_gene | testis_specific | FALSE | 6 |
| 27 | ENSG00000251258 | RFPL4B | CT_gene | testis_specific | TRUE | 6 |
| 28 | ENSG00000230533 | LINC03004 | CT_gene | testis_specific | FALSE | 6 |
| 29 | ENSG00000164744 | SUN3 | CT_gene | testis_specific | TRUE | 7 |
| 30 | ENSG00000234707 | SEC61G-DT | CT_gene | testis_specific | FALSE | 7 |
| 31 | ENSG00000241149 | ZNF722 | CT_gene | testis_specific | FALSE | 7 |
| 32 | ENSG00000232729 | GTF2I-AS1 | CT_gene | testis_specific | FALSE | 7 |
| 33 | ENSG00000176566 | DCAF4L2 | CT_gene | testis_specific | TRUE | 8 |
| 34 | ENSG00000137090 | DMRT1 | CT_gene | testis_specific | FALSE | 9 |
| 35 | ENSG00000177910 | SPATA31C2 | CT_gene | testis_specific | TRUE | 9 |
| 36 | ENSG00000230601 | TEX48 | CT_gene | testis_specific | TRUE | 9 |
| 37 | ENSG00000229205 | LINC00200 | CT_gene | testis_specific | TRUE | 10 |
| 38 | ENSG00000228636 | LINC02663 | CT_gene | testis_specific | TRUE | 10 |
| 39 | ENSG00000233515 | LINC01518 | CT_gene | testis_specific | TRUE | 10 |
| 40 | ENSG00000254518 | LINC02750 | CT_gene | testis_specific | FALSE | 11 |
| 41 | ENSG00000168070 | MAJIN | CT_gene | testis_specific | FALSE | 11 |
| 42 | ENSG00000249196 | TMEM132D-AS1 | CT_gene | testis_specific | TRUE | 12 |
| 43 | ENSG00000279516 | FAM230C | CT_gene | testis_specific | TRUE | 13 |
| 44 | ENSG00000198033 | TUBA3C | CT_gene | testis_specific | TRUE | 13 |
| 45 | ENSG00000132972 | RNF17 | CT_gene | testis_specific | TRUE | 13 |
| 46 | ENSG00000174015 | CBY2 | CT_gene | testis_specific | FALSE | 13 |
| 47 | ENSG00000172717 | GARIN2 | CT_gene | testis_specific | FALSE | 14 |
| 48 | ENSG00000258710 | LINC01193 | CT_gene | testis_specific | FALSE | 15 |
| 49 | ENSG00000261649 | GOLGA6L7 | CT_gene | testis_specific | TRUE | 15 |
| 50 | ENSG00000260172 | LINC01413 | CT_gene | testis_specific | TRUE | 15 |
| 51 | ENSG00000162039 | MEIOB | CT_gene | testis_specific | FALSE | 16 |
| 52 | ENSG00000126856 | PRDM7 | CT_gene | testis_specific | FALSE | 16 |
| 53 | ENSG00000223510 | CDRT15 | CT_gene | testis_specific | FALSE | 17 |
| 54 | ENSG00000276399 | FLJ36000 | CT_gene | testis_specific | TRUE | 17 |
| 55 | ENSG00000141371 | CHCT1 | CT_gene | testis_specific | FALSE | 17 |
| 56 | ENSG00000132204 | LINC00470 | CT_gene | testis_specific | TRUE | 18 |
| 57 | ENSG00000183206 | POTEC | CT_gene | testis_specific | TRUE | 18 |
| 58 | ENSG00000263711 | LINC02864 | CT_gene | testis_specific | TRUE | 18 |
| 59 | ENSG00000196350 | ZNF729 | CT_gene | testis_specific | FALSE | 19 |
| 60 | ENSG00000268696 | ZNF723 | CT_gene | testis_specific | FALSE | 19 |
| 61 | ENSG00000261949 | GFY | CT_gene | testis_specific | TRUE | 19 |
| 62 | ENSG00000160505 | NLRP4 | CT_gene | testis_specific | TRUE | 19 |
| 63 | ENSG00000244588 | RAD21L1 | CT_gene | testis_specific | FALSE | 20 |
| 64 | ENSG00000125823 | CSTL1 | CT_gene | testis_specific | FALSE | 20 |
| 65 | ENSG00000124092 | CTCFL | CT_gene | testis_specific | TRUE | 20 |
| 66 | ENSG00000198054 | DSCR8 | CT_gene | testis_specific | TRUE | 21 |
| 67 | ENSG00000169059 | VCX3A | CT_gene | testis_specific | TRUE | X |
| 68 | ENSG00000182583 | VCX | CT_gene | testis_specific | TRUE | X |
| 69 | ENSG00000177504 | VCX2 | CT_gene | testis_specific | TRUE | X |
| 70 | ENSG00000205642 | VCX3B | CT_gene | testis_specific | TRUE | X |
| 71 | ENSG00000184735 | DDX53 | CT_gene | testis_specific | TRUE | X |
| 72 | ENSG00000176746 | MAGEB6 | CT_gene | testis_specific | TRUE | X |

| | | | | | | |
|---|---|---|---|---|---|---|
| 73 | ENSG00000224960 | PPP4R3C | CT_gene | testis_specific | TRUE | X |
| 74 | ENSG00000189186 | DCAF8L2 | CT_gene | testis_specific | TRUE | X |
| 75 | ENSG00000099399 | MAGEB2 | CT_gene | testis_specific | TRUE | X |
| 76 | ENSG00000214107 | MAGEB1 | CT_gene | testis_specific | TRUE | X |
| 77 | ENSG00000132446 | FTHL17 | CT_gene | testis_specific | TRUE | X |
| 78 | ENSG00000189023 | MAGEB16 | CT_gene | testis_specific | TRUE | X |
| 79 | ENSG00000165583 | SSX5 | CT_gene | testis_specific | TRUE | X |
| 80 | ENSG00000126752 | SSX1 | CT_gene | testis_specific | TRUE | X |
| 81 | ENSG00000274274 | GAGE13 | CT_gene | testis_specific | TRUE | X |
| 82 | ENSG00000216649 | GAGE12E | CT_gene | testis_specific | TRUE | X |
| 83 | ENSG00000205777 | GAGE1 | CT_gene | testis_specific | TRUE | X |
| 84 | ENSG00000189064 | GAGE2A | CT_gene | testis_specific | TRUE | X |
| 85 | ENSG00000068985 | PAGE1 | CT_gene | testis_specific | TRUE | X |
| 86 | ENSG00000283093 | CENPVL2 | CT_gene | testis_specific | TRUE | X |
| 87 | ENSG00000223591 | CENPVL1 | CT_gene | testis_specific | TRUE | X |
| 88 | ENSG00000204379 | XAGE1A | CT_gene | testis_specific | TRUE | X |
| 89 | ENSG00000204382 | XAGE1B | CT_gene | testis_specific | TRUE | X |
| 90 | ENSG00000241476 | SSX2 | CT_gene | testis_specific | TRUE | X |
| 91 | ENSG00000268447 | SSX2B | CT_gene | testis_specific | TRUE | X |
| 92 | ENSG00000171405 | XAGE5 | CT_gene | testis_specific | TRUE | X |
| 93 | ENSG00000234068 | PAGE2 | CT_gene | testis_specific | TRUE | X |
| 94 | ENSG00000158639 | PAGE5 | CT_gene | testis_specific | TRUE | X |
| 95 | ENSG00000215115 | CXorf49 | CT_gene | testis_specific | TRUE | X |
| 96 | ENSG00000215113 | CXorf49B | CT_gene | testis_specific | TRUE | X |
| 97 | ENSG00000153779 | TGIF2LX | CT_gene | testis_specific | TRUE | X |
| 98 | ENSG00000269405 | NXF2 | CT_gene | testis_specific | TRUE | X |
| 99 | ENSG00000123576 | ESX1 | CT_gene | testis_specific | FALSE | X |
| 100 | ENSG00000226685 | CT47A12 | CT_gene | testis_specific | TRUE | X |
| 101 | ENSG00000226929 | CT47A11 | CT_gene | testis_specific | TRUE | X |
| 102 | ENSG00000236371 | CT47A1 | CT_gene | testis_specific | TRUE | X |
| 103 | ENSG00000282815 | TEX13C | CT_gene | testis_specific | TRUE | X |
| 104 | ENSG00000169551 | CT55 | CT_gene | testis_specific | TRUE | X |
| 105 | ENSG00000268940 | CT45A1 | CT_gene | testis_specific | TRUE | X |
| 106 | ENSG00000269096 | CT45A3 | CT_gene | testis_specific | TRUE | X |
| 107 | ENSG00000228836 | CT45A5 | CT_gene | testis_specific | TRUE | X |
| 108 | ENSG00000278289 | CT45A6 | CT_gene | testis_specific | TRUE | X |
| 109 | ENSG00000273696 | CT45A7 | CT_gene | testis_specific | TRUE | X |
| 110 | ENSG00000278085 | CT45A8 | CT_gene | testis_specific | TRUE | X |
| 111 | ENSG00000270946 | CT45A9 | CT_gene | testis_specific | TRUE | X |
| 112 | ENSG00000269586 | CT45A10 | CT_gene | testis_specific | TRUE | X |
| 113 | ENSG00000181433 | SAGE1 | CT_gene | testis_specific | TRUE | X |
| 114 | ENSG00000227234 | SPANXB1 | CT_gene | testis_specific | TRUE | X |
| 115 | ENSG00000198573 | SPANXC | CT_gene | testis_specific | TRUE | X |
| 116 | ENSG00000196406 | SPANXD | CT_gene | testis_specific | TRUE | X |
| 117 | ENSG00000155495 | MAGEC1 | CT_gene | testis_specific | TRUE | X |
| 118 | ENSG00000046774 | MAGEC2 | CT_gene | testis_specific | TRUE | X |
| 119 | ENSG00000235699 | CXorf51B | CT_gene | testis_specific | TRUE | X |
| 120 | ENSG00000176988 | FMR1NB | CT_gene | testis_specific | TRUE | X |
| 121 | ENSG00000267978 | MAGEA9B | CT_gene | testis_specific | TRUE | X |
| 122 | ENSG00000185247 | MAGEA11 | CT_gene | testis_specific | TRUE | X |
| 123 | ENSG00000123584 | MAGEA9 | CT_gene | testis_specific | TRUE | X |
| 124 | ENSG00000156009 | MAGEA8 | CT_gene | testis_specific | TRUE | X |
| 125 | ENSG00000166049 | PASD1 | CT_gene | testis_specific | TRUE | X |
| 126 | ENSG00000229967 | MAGEA4-AS1 | CT_gene | testis_specific | TRUE | X |
| 127 | ENSG00000147381 | MAGEA4 | CT_gene | testis_specific | TRUE | X |
| 128 | ENSG00000124260 | MAGEA10 | CT_gene | testis_specific | TRUE | X |
| 129 | ENSG00000221867 | MAGEA3 | CT_gene | testis_specific | TRUE | X |
| 130 | ENSG00000213401 | MAGEA12 | CT_gene | testis_specific | TRUE | X |
| 131 | ENSG00000268606 | MAGEA2 | CT_gene | testis_specific | TRUE | X |
| 132 | ENSG00000197172 | MAGEA6 | CT_gene | testis_specific | TRUE | X |
| 133 | ENSG00000198681 | MAGEA1 | CT_gene | testis_specific | TRUE | X |
| 134 | ENSG00000268651 | CTAG1A | CT_gene | testis_specific | TRUE | X |
| 135 | ENSG00000184033 | CTAG1B | CT_gene | testis_specific | TRUE | X |
| 136 | ENSG00000126890 | CTAG2 | CT_gene | testis_specific | TRUE | X |
| 137 | ENSG00000168757 | TSPY2 | CT_gene | testis_specific | FALSE | Y |
| 138 | ENSG00000233803 | TSPY4 | CT_gene | testis_specific | FALSE | Y |
| 139 | ENSG00000228927 | TSPY3 | CT_gene | testis_specific | FALSE | Y |
| 140 | ENSG00000258992 | TSPY1 | CT_gene | testis_specific | FALSE | Y |
| 141 | ENSG00000238074 | TSPY9 | CT_gene | testis_specific | FALSE | Y |
| 142 | ENSG00000129864 | VCY | CT_gene | testis_specific | TRUE | Y |
| 143 | ENSG00000129862 | VCY1B | CT_gene | testis_specific | TRUE | Y |
| 144 | ENSG00000242875 | RBMY1B | CT_gene | testis_specific | TRUE | Y |
| 145 | ENSG00000234414 | RBMY1A1 | CT_gene | testis_specific | TRUE | Y |
| 146 | ENSG00000242389 | RBMY1E | CT_gene | testis_specific | TRUE | Y |

**Table S2: List of 134 CG-Preferential genes**

| | ensembl_gene_id | external_gene_name | CT_gene_type | testis_specificity | regulated_by_methylation | chr |
|---|---|---|---|---|---|---|
| 1 | ENSG00000162571 | TTLL10 | CTP_gene | testis_preferential | TRUE | 1 |
| 2 | ENSG00000236423 | LINC01134 | CTP_gene | testis_preferential | FALSE | 1 |
| 3 | ENSG00000233421 | LINC01783 | CTP_gene | testis_preferential | FALSE | 1 |
| 4 | ENSG00000233203 | DHCR24-DT | CTP_gene | testis_preferential | FALSE | 1 |
| 5 | ENSG00000226088 | LINC03102 | CTP_gene | testis_preferential | TRUE | 1 |
| 6 | ENSG00000162620 | LRRIQ3 | CTP_gene | testis_preferential | FALSE | 1 |
| 7 | ENSG00000162641 | AKNAD1 | CTP_gene | testis_preferential | TRUE | 1 |
| 8 | ENSG00000215866 | LINC01356 | CTP_gene | testis_preferential | FALSE | 1 |
| 9 | ENSG00000143452 | HORMAD1 | CTP_gene | testis_preferential | TRUE | 1 |
| 10 | ENSG00000160838 | LRRC71 | CTP_gene | testis_preferential | FALSE | 1 |
| 11 | ENSG00000171722 | SPATA46 | CTP_gene | testis_preferential | TRUE | 1 |
| 12 | ENSG00000143194 | MAEL | CTP_gene | testis_preferential | TRUE | 1 |
| 13 | ENSG00000162779 | AXDND1 | CTP_gene | testis_preferential | FALSE | 1 |
| 14 | ENSG00000186007 | LEMD1 | CTP_gene | testis_preferential | FALSE | 1 |
| 15 | ENSG00000162814 | SPATA17 | CTP_gene | testis_preferential | FALSE | 1 |
| 16 | ENSG00000173728 | SPMIP3 | CTP_gene | testis_preferential | TRUE | 1 |
| 17 | ENSG00000163793 | DNAJC5G | CTP_gene | testis_preferential | FALSE | 2 |
| 18 | ENSG00000271889 | C2orf74-AS1 | CTP_gene | testis_preferential | FALSE | 2 |
| 19 | ENSG00000234199 | LINC01191 | CTP_gene | testis_preferential | TRUE | 2 |
| 20 | ENSG00000152086 | TUBA3E | CTP_gene | testis_preferential | TRUE | 2 |
| 21 | ENSG00000188738 | FSIP2 | CTP_gene | testis_preferential | FALSE | 2 |
| 22 | ENSG00000155754 | C2CD6 | CTP_gene | testis_preferential | FALSE | 2 |
| 23 | ENSG00000242242 | NECTIN3-AS1 | CTP_gene | testis_preferential | FALSE | 3 |
| 24 | ENSG00000065371 | ROPN1 | CTP_gene | testis_preferential | FALSE | 3 |
| 25 | ENSG00000114547 | ROPN1B | CTP_gene | testis_preferential | FALSE | 3 |
| 26 | ENSG00000114656 | CFAP92 | CTP_gene | testis_preferential | FALSE | 3 |
| 27 | ENSG00000227375 | DLG1-AS1 | CTP_gene | testis_preferential | FALSE | 3 |
| 28 | ENSG00000246095 | LINC01096 | CTP_gene | testis_preferential | FALSE | 4 |
| 29 | ENSG00000182308 | DCAF4L1 | CTP_gene | testis_preferential | TRUE | 4 |
| 30 | ENSG00000156269 | NAA11 | CTP_gene | testis_preferential | TRUE | 4 |
| 31 | ENSG00000137473 | TTC29 | CTP_gene | testis_preferential | FALSE | 4 |
| 32 | ENSG00000150628 | SPATA4 | CTP_gene | testis_preferential | FALSE | 4 |
| 33 | ENSG00000251221 | LINC01337 | CTP_gene | testis_preferential | FALSE | 5 |
| 34 | ENSG00000153347 | FAM81B | CTP_gene | testis_preferential | FALSE | 5 |
| 35 | ENSG00000223442 | TH2LCRR | CTP_gene | testis_preferential | FALSE | 5 |
| 36 | ENSG00000249346 | LINC01016 | CTP_gene | testis_preferential | FALSE | 6 |
| 37 | ENSG00000204091 | TDRG1 | CTP_gene | testis_preferential | TRUE | 6 |
| 38 | ENSG00000124490 | CRISP2 | CTP_gene | testis_preferential | FALSE | 6 |
| 39 | ENSG00000203907 | OOEP | CTP_gene | testis_preferential | TRUE | 6 |
| 40 | ENSG00000203711 | LINC02901 | CTP_gene | testis_preferential | FALSE | 6 |
| 41 | ENSG00000146453 | PNLDC1 | CTP_gene | testis_preferential | TRUE | 6 |
| 42 | ENSG00000196335 | STK31 | CTP_gene | testis_preferential | TRUE | 7 |
| 43 | ENSG00000175877 | TMEM270 | CTP_gene | testis_preferential | FALSE | 7 |
| 44 | ENSG00000242950 | ERVW-1 | CTP_gene | testis_preferential | TRUE | 7 |
| 45 | ENSG00000106013 | ANKRD7 | CTP_gene | testis_preferential | FALSE | 7 |
| 46 | ENSG00000235631 | RNF148 | CTP_gene | testis_preferential | FALSE | 7 |
| 47 | ENSG00000135248 | GARIN1B | CTP_gene | testis_preferential | TRUE | 7 |
| 48 | ENSG00000158525 | CPA5 | CTP_gene | testis_preferential | TRUE | 7 |
| 49 | ENSG00000165131 | LLCFC1 | CTP_gene | testis_preferential | FALSE | 7 |
| 50 | ENSG00000239377 | ZNF775-AS1 | CTP_gene | testis_preferential | FALSE | 7 |
| 51 | ENSG00000164900 | GBX1 | CTP_gene | testis_preferential | FALSE | 7 |
| 52 | ENSG00000248538 | PPP1R3B-DT | CTP_gene | testis_preferential | TRUE | 8 |
| 53 | ENSG00000147570 | DNAJC5B | CTP_gene | testis_preferential | FALSE | 8 |
| 54 | ENSG00000245080 | MIR3150BHG | CTP_gene | testis_preferential | FALSE | 8 |
| 55 | ENSG00000186583 | SPATC1 | CTP_gene | testis_preferential | TRUE | 8 |
| 56 | ENSG00000185972 | CCIN | CTP_gene | testis_preferential | TRUE | 9 |
| 57 | ENSG00000225684 | FAM225B | CTP_gene | testis_preferential | FALSE | 9 |
| 58 | ENSG00000231528 | FAM225A | CTP_gene | testis_preferential | FALSE | 9 |
| 59 | ENSG00000167094 | TTC16 | CTP_gene | testis_preferential | FALSE | 9 |
| 60 | ENSG00000185863 | TMEM210 | CTP_gene | testis_preferential | FALSE | 9 |
| 61 | ENSG00000165383 | LRRC18 | CTP_gene | testis_preferential | FALSE | 10 |
| 62 | ENSG00000222047 | C10orf55 | CTP_gene | testis_preferential | FALSE | 10 |
| 63 | ENSG00000166796 | LDHC | CTP_gene | testis_preferential | FALSE | 11 |
| 64 | ENSG00000255236 | CHKA-DT | CTP_gene | testis_preferential | FALSE | 11 |
| 65 | ENSG00000255507 | UVRAG-DT | CTP_gene | testis_preferential | FALSE | 11 |
| 66 | ENSG00000178795 | GDPD4 | CTP_gene | testis_preferential | TRUE | 11 |

| | | | | | | | |
|---|---|---|---|---|---|---|---|
| 67 | ENSG00000165325 | DEUP1 | CTP_gene | testis_preferential | FALSE | | 11 |
| 68 | ENSG00000173262 | SLC2A14 | CTP_gene | testis_preferential | FALSE | | 12 |
| 69 | ENSG00000111305 | GSG1 | CTP_gene | testis_preferential | FALSE | | 12 |
| 70 | ENSG00000170627 | GTSF1 | CTP_gene | testis_preferential | TRUE | | 12 |
| 71 | ENSG00000139351 | SYCP3 | CTP_gene | testis_preferential | TRUE | | 12 |
| 72 | ENSG00000179088 | C12orf42 | CTP_gene | testis_preferential | FALSE | | 12 |
| 73 | ENSG00000151846 | PABPC3 | CTP_gene | testis_preferential | FALSE | | 13 |
| 74 | ENSG00000120669 | SOHLH2 | CTP_gene | testis_preferential | TRUE | | 13 |
| 75 | ENSG00000242715 | CCDC169 | CTP_gene | testis_preferential | FALSE | | 13 |
| 76 | ENSG00000231817 | LINC01198 | CTP_gene | testis_preferential | FALSE | | 13 |
| 77 | ENSG00000231473 | RB1-DT | CTP_gene | testis_preferential | FALSE | | 13 |
| 78 | ENSG00000165496 | RPL10L | CTP_gene | testis_preferential | TRUE | | 14 |
| 79 | ENSG00000196553 | CCDC196 | CTP_gene | testis_preferential | FALSE | | 14 |
| 80 | ENSG00000258584 | FAM181A-AS1 | CTP_gene | testis_preferential | FALSE | | 14 |
| 81 | ENSG00000212766 | EWSAT1 | CTP_gene | testis_preferential | FALSE | | 15 |
| 82 | ENSG00000225362 | CT62 | CTP_gene | testis_preferential | FALSE | | 15 |
| 83 | ENSG00000260469 | INSYN1-AS1 | CTP_gene | testis_preferential | FALSE | | 15 |
| 84 | ENSG00000117971 | CHRNB4 | CTP_gene | testis_preferential | FALSE | | 15 |
| 85 | ENSG00000261441 | POLG-DT | CTP_gene | testis_preferential | FALSE | | 15 |
| 86 | ENSG00000153060 | TEKT5 | CTP_gene | testis_preferential | FALSE | | 16 |
| 87 | ENSG00000168676 | KCTD19 | CTP_gene | testis_preferential | FALSE | | 16 |
| 88 | ENSG00000159708 | LRRC36 | CTP_gene | testis_preferential | FALSE | | 16 |
| 89 | ENSG00000141096 | DPEP3 | CTP_gene | testis_preferential | TRUE | | 16 |
| 90 | ENSG00000141255 | SPATA22 | CTP_gene | testis_preferential | TRUE | | 17 |
| 91 | ENSG00000177294 | FBXO39 | CTP_gene | testis_preferential | FALSE | | 17 |
| 92 | ENSG00000171931 | FBXW10 | CTP_gene | testis_preferential | FALSE | | 17 |
| 93 | ENSG00000141316 | SPACA3 | CTP_gene | testis_preferential | FALSE | | 17 |
| 94 | ENSG00000270806 | C17orf50 | CTP_gene | testis_preferential | FALSE | | 17 |
| 95 | ENSG00000186075 | ZPBP2 | CTP_gene | testis_preferential | FALSE | | 17 |
| 96 | ENSG00000180336 | MEIOC | CTP_gene | testis_preferential | FALSE | | 17 |
| 97 | ENSG00000121101 | TEX14 | CTP_gene | testis_preferential | FALSE | | 17 |
| 98 | ENSG00000259349 | APPBP2-DT | CTP_gene | testis_preferential | FALSE | | 17 |
| 99 | ENSG00000173838 | MARCHF10 | CTP_gene | testis_preferential | FALSE | | 17 |
| 100 | ENSG00000178404 | CEP295NL | CTP_gene | testis_preferential | FALSE | | 17 |
| 101 | ENSG00000265692 | LINC01970 | CTP_gene | testis_preferential | FALSE | | 17 |
| 102 | ENSG00000182459 | TEX19 | CTP_gene | testis_preferential | FALSE | | 17 |
| 103 | ENSG00000173213 | TUBB8B | CTP_gene | testis_preferential | TRUE | | 18 |
| 104 | ENSG00000265933 | LINC00668 | CTP_gene | testis_preferential | TRUE | | 18 |
| 105 | ENSG00000154611 | PSMA8 | CTP_gene | testis_preferential | TRUE | | 18 |
| 106 | ENSG00000105549 | SPMAP2 | CTP_gene | testis_preferential | TRUE | | 19 |
| 107 | ENSG00000198723 | SAXO5 | CTP_gene | testis_preferential | FALSE | | 19 |
| 108 | ENSG00000198028 | ZNF560 | CTP_gene | testis_preferential | FALSE | | 19 |
| 109 | ENSG00000173809 | TDRD12 | CTP_gene | testis_preferential | TRUE | | 19 |
| 110 | ENSG00000105679 | GAPDHS | CTP_gene | testis_preferential | FALSE | | 19 |
| 111 | ENSG00000131126 | TEX101 | CTP_gene | testis_preferential | TRUE | | 19 |
| 112 | ENSG00000105467 | SYNGR4 | CTP_gene | testis_preferential | FALSE | | 19 |
| 113 | ENSG00000104901 | DKKL1 | CTP_gene | testis_preferential | FALSE | | 19 |
| 114 | ENSG00000258713 | C20orf141 | CTP_gene | testis_preferential | FALSE | | 20 |
| 115 | ENSG00000215529 | EFCAB8 | CTP_gene | testis_preferential | FALSE | | 20 |
| 116 | ENSG00000124196 | GTSF1L | CTP_gene | testis_preferential | FALSE | | 20 |
| 117 | ENSG00000237232 | ZNF295-AS1 | CTP_gene | testis_preferential | FALSE | | 21 |
| 118 | ENSG00000185686 | PRAME | CTP_gene | testis_preferential | TRUE | | 22 |
| 119 | ENSG00000100121 | GGTLC2 | CTP_gene | testis_preferential | FALSE | | 22 |
| 120 | ENSG00000178248 | FAM230I | CTP_gene | testis_preferential | FALSE | | 22 |
| 121 | ENSG00000128322 | IGLL1 | CTP_gene | testis_preferential | FALSE | | 22 |
| 122 | ENSG00000235989 | MORC2-AS1 | CTP_gene | testis_preferential | FALSE | | 22 |
| 123 | ENSG00000086717 | PPEF1 | CTP_gene | testis_preferential | FALSE | X | |
| 124 | ENSG00000224659 | GAGE12J | CTP_gene | testis_preferential | TRUE | X | |
| 125 | ENSG00000275113 | GAGE2E | CTP_gene | testis_preferential | TRUE | X | |
| 126 | ENSG00000237671 | GAGE12C | CTP_gene | testis_preferential | TRUE | X | |
| 127 | ENSG00000236362 | GAGE12F | CTP_gene | testis_preferential | TRUE | X | |
| 128 | ENSG00000224902 | GAGE12H | CTP_gene | testis_preferential | TRUE | X | |
| 129 | ENSG00000147082 | CCNB3 | CTP_gene | testis_preferential | FALSE | X | |
| 130 | ENSG00000238269 | PAGE2B | CTP_gene | testis_preferential | TRUE | X | |
| 131 | ENSG00000102387 | TAF7L | CTP_gene | testis_preferential | FALSE | X | |
| 132 | ENSG00000204019 | CT83 | CTP_gene | testis_preferential | TRUE | X | |
| 133 | ENSG00000186471 | AKAP14 | CTP_gene | testis_preferential | FALSE | X | |
| 134 | ENSG00000007350 | TKTL1 | CTP_gene | testis_preferential | TRUE | X | |

**Table S3.** List of datasets used in CTexploreR analyses

| Type of data | Source | Accession code / File |
|---|---|---|
| RNASeq fastq files from normal tissues | ENCODE [77] | testis (ENCFF140UYT, ENCFF794EAB)<br>thyroid gland (ENCFF151GUG, ENCFF628TMU)<br>gastrocnemius medialis (ENCFF004CNM, ENCFF086LCO)<br>adrenal gland (ENCFF911BTP, ENCFF904VHM)<br>subcutaneous adipose tissue (ENCFF667CWY, ENCFF360KZB)<br>stomach (ENCFF741NGG, ENCFF582ILA)<br>upper lobe of left lung (ENCFF719YBM, ENCFF801ZKX)<br>suprapubic skin (ENCFF398KGB, ENCFF058JYK)<br>breast epithelium (ENCFF767QVV, ENCFF050GYP)<br>lower leg skin (ENCFF431RAQ, ENCFF008OVI)<br>sigmoid colon (ENCFF153ULW, ENCFF182OWD)<br>transverse colon (ENCFF411UIT, ENCFF992NAN)<br>spleen (ENCFF567ORO, ENCFF743QDT)<br>gastroesophageal sphincter (ENCFF250BPC, ENCFF842DCO)<br>tibial nerve (ENCFF534AYT, ENCFF935LBC)<br>esophagus muscularis mucosa (ENCFF091UZU, ENCFF163DLM)<br>omental fat pad (ENCFF745PTG, ENCFF824ZLA)<br>esophagus squamous epithelium (ENCFF585TOW, ENCFF230GLF) |
| RNA-Seq from cell lines treated with 5-Aza | ENCODE[77] | IMR5-75 CTL (SRR3326020/SRR3326021)<br>IMR5-75 DAC (SRR3326022/SRR3326023)<br>HCT116 CTL (SRR9108737/SRR9108738)<br>HCT116 DAC (SRR9108739/SRR9108740)<br>HEK293T CTL (SRR1618781/SRR1618782)<br>HEK293T DAC (SRR1618783/SRR1618784)<br>HMLER DAC (SRR3362409/SRR3362410)<br>HMLER CTL (SRR3362411/SRR3362412)<br>NCH612 CTL (SRR12105788/SRR12105789)<br>NCH612 DAC (SRR12105790/SRR12105791)<br>NCH1681 CTL (SRR12105792/SRR12105793)<br>NCH1681 DAC (SRR12105794/SRR12105795)<br>TS603 CTL (SRR12105780/SRR12105781)<br>TS603 DAC (SRR12105782/SRR12105783)<br>B2-1 CTL (SRR5363797/SRR5363798)<br>B2-1 DAC (SRR5363799/SRR5363800) |
| WGBS from healthy tissues | ENCODE[77] | adipose ENCFF318AMC<br>colon ENCFF157POM<br>oesophagus ENCFF625GVK<br>heart ENCFF536RSX<br>intestine ENCFF241AQC<br>lung ENCFF039JFT<br>muscle ENCFF121ZES<br>pancreas ENCFF763RUE<br>placenta ENCFF437OKM<br>skin ENCFF219GCQ<br>stomach ENCFF497YOO<br>testis ENCFF715DMX<br>thyroid ENCFF223LJW |
